# *Shigella* ipaA mediates actin bundling through diffusible vinculin oligomers with activation imprint

**DOI:** 10.1101/2022.11.07.515412

**Authors:** Cesar Valencia-Gallardo, Daniel-Isui Aguilar-Salvador, Hamed Khakzad, Benjamin Cocom-Chan, Charles Bou-Nader, Christophe Velours, Yosra Zarrouk, Christophe Le Clainche, Christian Malosse, Diogo Borges Lima, Nicole Quenech’Du, Bilal Mazhar, Sami Essid, Marc Fontecave, Atef Asnacios, Julia Chamot-Rooke, Lars Malmström, Guy Tran Van Nhieu

## Abstract

Upon activation, vinculin reinforces cytoskeletal anchorage during cell adhesion. Activating ligands classically disrupt intramolecular interactions between the vinculin head and tail domain that binds to actin filaments. Here, we show that *Shigella* IpaA triggers major allosteric changes in the head domain leading to vinculin homo-oligomerization. Through the cooperative binding of its three vinculin-binding sites (VBSs), IpaA induces a striking reorientation of the D1 and D2 head subdomains associated with vinculin oligomerization. IpaA thus acts as a catalyst producing vinculin clusters that bundle actin at a distance from the activation site and trigger the formation of highly stable adhesions resisting the action of actin relaxing drugs. Unlike canonical activation, vinculin homo-oligomers induced by IpaA appear to keep a persistent imprint of the activated state in addition to their bundling activity, accounting for stable cell adhesion independent of force transduction and relevant to bacterial invasion.

## Introduction

*Shigella*, the causative agent of bacillary dysentery, invades epithelial cells by injecting type III effectors that locally reorganize the actin cytoskeleton (Ogawa, Handa et al. 2008, Valencia-Gallardo, Carayol et al. 2015, Mattock and Blocker 2017). *Shigella* invasion involves limited contacts with host cells and critically depends on the type III effector IpaA that promotes cytoskeletal anchorage by targeting the (FA) adhesion proteins talin, and vinculin (Romero, Grompone et al. 2011, Valencia-Gallardo, Carayol et al. 2015, Valencia-Gallardo, Bou-Nader et al. 2019). During integrin-mediated cell adhesion, talin acts as a mechanosensor by exposing its vinculin binding sites (VBSs) that recruit and activate vinculin, reinforcing anchorage to the actin cytoskeleton in response to mechanical load (Humphries, Wang et al. 2007, Parsons, Horwitz et al. 2010, Lavelin, Wolfenson et al. 2013, Ciobanasu, Faivre and Le Clainche 2014, Atherton, Stutchbury et al. 2015). *Shigella* cannot generate the type of mechanical load required for strong cytoskeletal anchorage, therefore, scaffolding of talin, and vinculin at bacterial invasion sites exclusively relies on IpaA. IpaA contains three vinculin binding sites (VBSs) in its carboxyterminal moiety, with diverse functions inferred from the crystal structures of complexes containing the VBS peptide (Izard, Tran Van Nhieu et al. 2006, Tran Van Nhieu and Izard 2007). Vinculin is classically described as a head domain (Vh) connected to a tail domain (Vt) by a flexible linker (Bakolitsa, Cohen et al. 2004). Vh contains three repetitions (D1-D3) of a conserved domain consisting of two antiparallel α-helix-bundles and a fourth α-helix bundle D4 (Bakolitsa, Cohen et al. 2004). A proline-rich unstructured linker bridges D4 and a five-helix bundle Vt containing the carboxyterminal F-actin binding domain (Bakolitsa, Cohen et al. 2004). Under its inactive folded form, intramolecular interactions between Vh and Vt prevent ligand binding. IpaA VBS1, as for all VBSs described to activate vinculin, interacts with the first helical bundle of the D1 domain, promoting major conformational changes that disrupt the Vh-Vt intramolecular interactions and free the vinculin F-actin binding region (Izard, Evans et al. 2004). IpaA VBS2, in contrast, interacts with the second helical bundle of D1 (Tran Van Nhieu and Izard 2007), hence, its association with IpaA VBS1 results in a very high affinity and stable IpaA VBS1-2:D1 complex, with an estimated K_D_ in the femtoM range (Tran Van Nhieu and Izard 2007). Functional evidence indicates that IpaA VBS3 cooperates with IpaA VBS1-2 to stimulate bacterial invasion (Park, Valencia-Gallardo et al. 2011, Valencia-Gallardo, Bou-Nader et al. 2019). IpaA VBS3, as an isolated peptide, acts as IpaA VBS1 by interacting with the vinculin D1 first helical bundle and promotes vinculin activation (Park, Valencia-Gallardo et al. 2011). IpaA VBS3, however, can also interact with talin to stimulate bacterial capture by filopodia during the early *Shigella* invasion phase of host cells (Valencia-Gallardo, Bou-Nader et al. 2019). The structural data indicate that IpaA VBS3 stabilizes the H1-H4 helix bundle expected to form in a partially stretched talin conformer at the low force range exerted by filopodia (Valencia-Gallardo, Bou-Nader et al. 2019). Intriguingly, IpaA VBS3 shares with talin VBS10 (H46) the ability to bind to vinculin and talin H1H4, suggesting a complex interplay between talin and vinculin during mechanotransduction (Valencia-Gallardo, Bou-Nader et al. 2019). Unlike talin VBSs, IpaA VBSs are not buried into helix bundles, presumably enabling targeting of vinculin and talin in a serendipitous manner.

In addition to strengthening cytoskeletal anchorage, vinculin has also been implicated in the bundling of actin filaments through dimerization via its tail domain (Vt), triggered by F-actin or phosphatidylinositol(4, 5) bisphosphate (PIP2) binding (Johnson and Craig 2000, Janssen, Kim et al. 2006, Chinthalapudi, Rangarajan et al. 2014). Consistent with a key role in Vt-mediated actin bundling, mutations in Vt that prevent PIP2-binding lead to defects in FA dynamics and formation (Chinthalapudi, Rangarajan et al. 2014). Also, mutations that prevent Vt dimerization or alter C-terminal hairpin involved in actin bundling lead to defects in FA formation and cell spreading, although the correlation between F-actin bundling activity and the amplitude of adhesion defects is unclear (Shen, Tolbert et al. 2011). The role of Vt-induced dimerization in scaffolding, however, remains unclear since it cannot simply explain the formation of high-order complexes. The formation of these high-order vinculin complexes could implicate the recruitment of other vinculin-binding partners or vinculin oligomerization mechanisms other than through Vt, possibly through Vh-Vh interactions observed in the so-called “parachute” structures (Molony and Burridge 1985). Of interest, upon activation, vinculin is known to promote the scaffolding of adhesion components during FA growth and maturation, a process that may also implicate its oligomerization (Thompson, Tolbert et al. 2013). More recently, through its interaction with branched actin networks and the bundling activity of talin-vinculin scaffolds, vinculin was also proposed to regulate the dynamics of actin polymerization at adhesion structures (Boujemaa-Paterski R, Martins B et al. 2020). These studies also pointed to the observation that vinculin-mediated actin bundling occurred at the site of vinculin activation (Boujemaa-Paterski R, Martins B et al. 2020), thereby imposing a frame spatially limiting the actin bundling activity of vinculin during adhesion maturation.

Here, we investigated the role of vinculin at the cell cortical sites of *Shigella* invasion, where all three IpaA VBSs are expected to bind target vinculin (Valencia-Gallardo, Bou-Nader et al. 2019). We show that the combined action of IpaA VBSs, induce major conformational changes in the vinculin head domain, a process that we coined “supra-activation”. These changes lead to the formation of vinculin homo-oligomers promoting the bundling of actin filaments at a distance from the activation site. Our results suggest that vinculin “supra-activation” also occurs during mechanotransduction and is required for maturation of cell adhesions.

## Results

### IpaA VBS1-3 are required for full recruitment of vinculin at *Shigella* contact sites

IpaA VBS3 targets a partially unfolded talin conformer, during early bacterial capture by filopodia, but is not expected to bind to fully activated talin (Valencia-Gallardo, Bou-Nader et al. 2019). Instead, at higher force ranges associated with FA maturation at the cell cortex, IpaA VBS3 is expected to target vinculin in combination with IpaA VBS1, 2. To test this, we analyzed adhesion structures induced by *Shigella* during bacterial invasion.

As shown in Fig. 1a and as previously reported (Tran Van Nhieu and Izard 2007), *Shigella* triggers the IpaA-dependent recruitment of vinculin at phagocytic cups (Fig. 1a, arrows, and 1b, WT). Vinculin recruitment was strongly reduced at bacterial contact sites induced by *ipaA/VBS3* expressing only IpaA VBS3 (Figs.1a, b, VBS3 and S1, Park, Valencia-Gallardo et al. 2011). *ipaA*/VBS3 triggered the recruitment of small vinculin patches but large phagocytic cups as observed for the wild-type strain were not detected (Figs. 1a, arrows and 1b). We previously identified mutations A495K and K498E in IpaA VBS3 affecting talin-but not vinculin-binding (Valencia-Gallardo, Bou-Nader et al. 2019). We found that these mutations did not affect the recruitment of vinculin small patches triggered by IpaA VBS3, consistent with a direct role of IpaA VBS3 in vinculin binding at bacterial contact sites (Figs. 1a, b, A495K and K498E; Fig. S1). Consistent with previous talin staining results, IpaA also induced the formation of vinculin-containing FAs distal to bacterial invasion sites (Fig. 1a, arrowheads; Valencia-Gallardo, Bou-Nader et al. 2019). In contrast to bacterial contact sites, these distal FAs formed at similar extents for WT *Shigella* and *ipaA*/VBS3 but were affected in talin-binding deficient VBS3 derivatives (Figs. 1a, arrowheads, 1c and S1), suggesting that vinculin recruitment at distal FAs occurred indirectly through talin.

**Figure 1.**
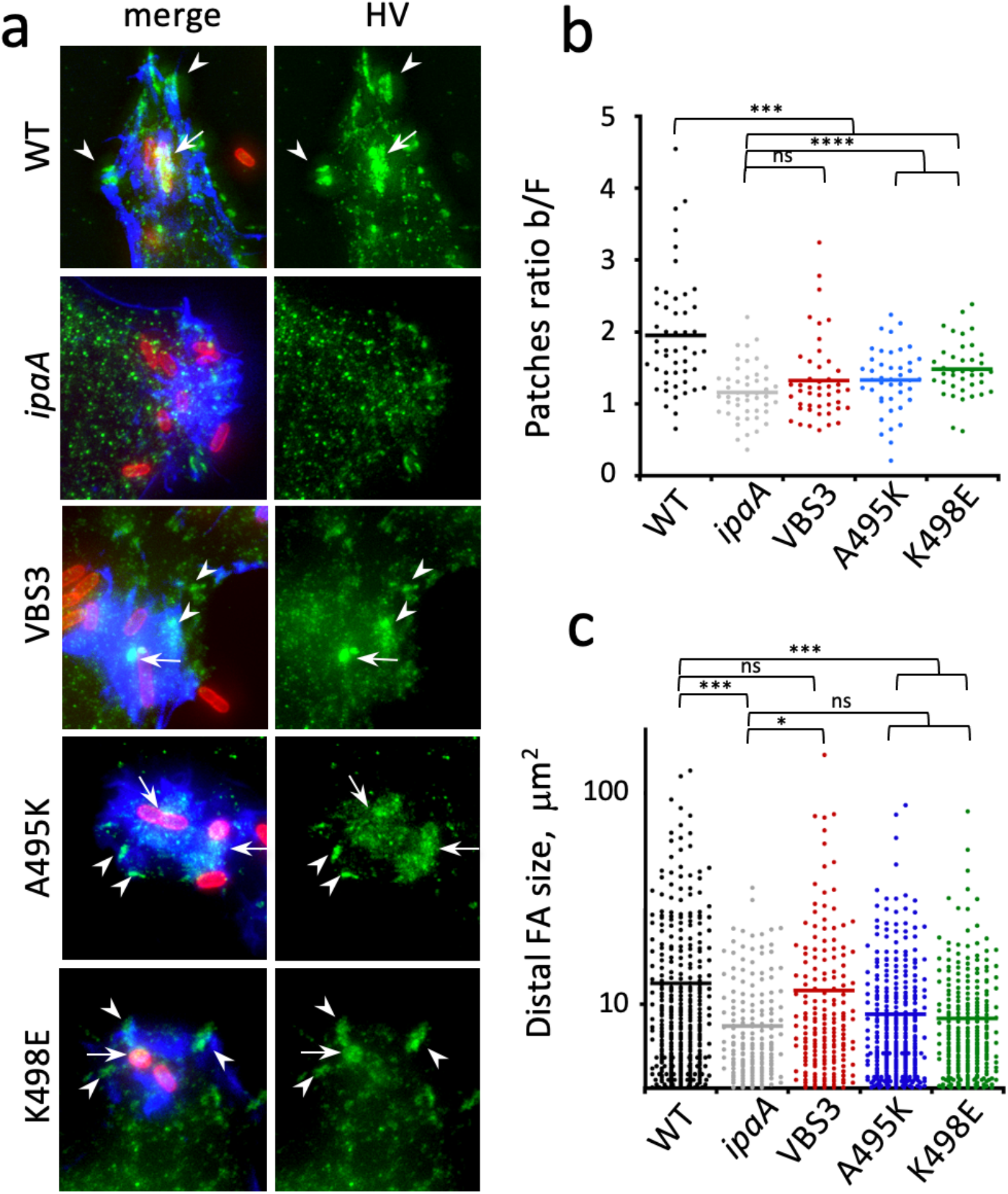
Vinculin recruitment at *Shigella* contact sites and distal adhesion structures during bacterial invasion. HeLa cells were challenged with bacteria for 30 min at 37°C, fixed and processed for immunofluorescence staining. *ipaA* mutant complemented with: full-length IpaA (WT); control vector (*ipaA*); IpaA ΔVBS1-2 (VBS3); IpaAΔVBS1-2 A495K (A495K); IpaAΔVBS1-2 K498E (K498E). a, representative micrographs. Merge: maximum projection of deconvolved confocal planes. vinculin: confocal plane corresponding to the cell basal surface. Red: bacteria; green: vinculin. Blue: F-actin. Vinculin recruitment at bacterial contact sites (arrows) and distal adhesion structures (arrowheads). Scale bar = 5 μm. b, vinculin recruitment at bacterial contact sites was quantified as the ratio of average fluorescence intensity of vinculin labeling associated with the bacterial body over that of actin foci (Star Methods, Fig. S1). The average ratio ± SEM is indicated. WT: 1.94 ± 0.11 (48 foci, N = 2); *ipaA*: 1.15 ± 0.05 (46 foci, N = 2); VBS3: 1.32± 0.08 (45 foci, N = 2); A495K: 1.33 ± 0.07 (42 foci, N = 2); K498E: 1.48 ± 0.06 (38 foci, N = 2). c, large vinculin adhesion structures were scored as detailed in the Star Methods section. Average FA size ± SEM μm^2^: WT: 12.54 ± 0.76 (393 FAs, N = 2); *ipaA*: 8.48 ± 0.49 (201 FAs, N = 2); VBS3: 11.59 ± 1.08 (207 FAs, N = 2); A495K: 8.96 ± 0.45 (376 FAs, N = 2); K498E: 8.56 ± 0.44 (291 FAs, N = 2). Mann and Whitney test: *: p < 0.05; ***: p <0.005; ****: p <0.001.

These results indicate that IpaA VBS3 binding to vinculin is required for the full recruitment of this cytoskeletal linker at phagocytic cups during *Shigella* invasion. In contrast, IpaA VBS3 appears to play a distinct role in bacterial-induced distal FAs, for which its talin-binding property is critical while vinculin binding is dispensable. These findings point at different functions of IpaA VBS3, contrasting with a mere role in vinculin scaffolding during *Shigella* invasion.

### IpaA induces vinculin higher order oligomerization

Previous analytical size exclusion chromatography (SEC) studies suggested that IpaA can bind to multiple vinculin molecules through its three VBSs (Park, Valencia-Gallardo et al. 2011). In the proposed model and akin to the model proposed for talin VBSs during mechanotransduction, each IpaA VBS binds to one vinculin molecule through interaction via the first bundle of the vinculin D1 subdomain, leading to its activation (Fig. 2a) (Park, Valencia-Gallardo et al. 2011). This view supports a redundant role for IpaA VBSs inconsistent with a differential role of IpaA VBS3 suggested in the previous set of experiments.

**Figure 2.**
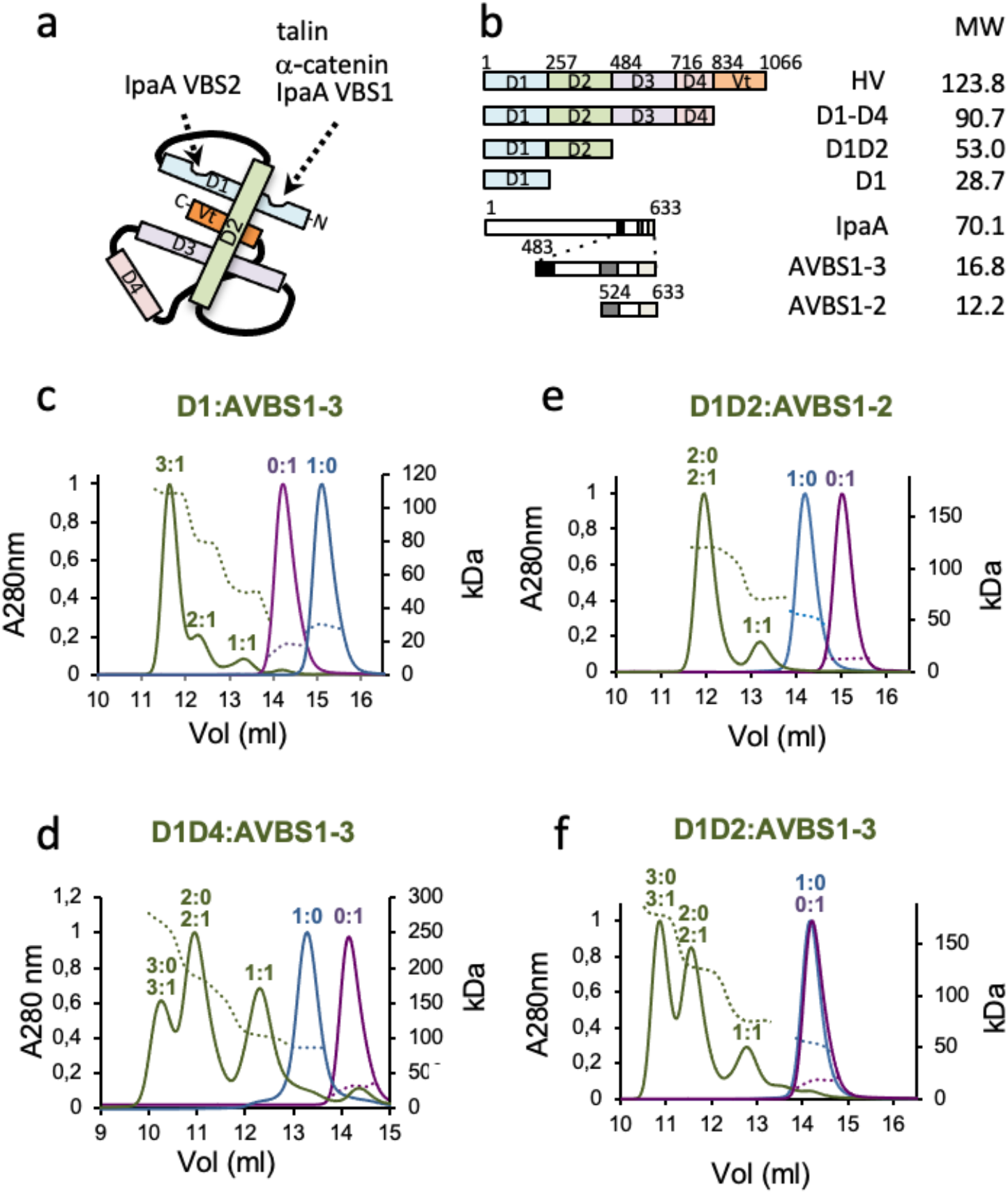
IpaA promotes vinculin homo-oligomerization. **a,** Scheme of folded vinculin (HV). The binding sites and corresponding ligands are indicated. **b,** Scheme of vinculin and IpaA constructs. vinculin domains and IpaA VBSs are depicted as boxes. The numbers indicate the start residue of each domain. MW: molecular weight in kDa. **c-f,** green: SEC elution profiles of complexes formed between AVBS1-3 (**c**, **d**, **f**) or AVBS1-2 (**e**) and the indicated vinculin derivatives; blue: the indicated vinculin derivative alone; purple: AVBS1-2 (**e**) or AVBS1-3 alone (**c**, **d**, **f**). The indicated complex stoichiometry was inferred from the molecular weight estimated by MALS. Dotted line: molecular weight.

To further characterize the role of IpaA VBS3 on vinculin binding, we studied the effects of the IpaA derivatives containing VBS1-2 (AVBS1-2) or VBS1-3 (AVBS1-3) on binding to derivatives containing different subdomains of the vinculin head (D1-D4) using SEC-MALS (Size Exclusion Chromatography-Multi-Angle Light Scattering) (Fig. 2b). When analyzing binding of AVBS1-3 to the D1 first subdomain of vinculin corresponding residues 1 −257 and consistent with previous SEC results (Park, Valencia-Gallardo et al. 2011), AVBS1-3:D1 complexes with a molar ration of 1:1, 2:1 and 3:1 were observed likely corresponding to the scaffolding of D1 molecules on the 3 IpaA VBSs based on the predicted molecular mass of the complexes (Fig. 2c). We then analyzed complexes formed upon incubation of AVBS1-3 with a construct containing vinculin residues 1-834 (D1D4), corresponding to full-length human vinculin (HV) devoid of the carboxyterminal F-actin binding domain (Fig. 2b). As shown in Fig. 2d, we found 1:1 D1D4:AVBS1-3 but unexpectedly, complexes containing 2 and 3 D1D4 molecules were also observed (Fig. 2d). Similar complexes containing 2 and 3 molecules of a derivative containing only the vinculin residues 1-484 (Fig. 2f, D1D2) upon incubation with AVBS1-3, indicating that vinculin oligomerization only required the vinculin D1 and D2 sub-domains. By contrast, when AVBS1-2 was incubated with D1D2, 1:1 and 2:0 - 2:1 D1D2:AVBS1-2 complexes were detected, but no D1D2 trimers (Fig. 2e). Because of the small size of AVBS1-2 and AVBS1-3 and the low extinction coefficient difference between complex partners, the detection limits of the SEC-MALS equipment did not allow to unambiguously distinguish between 2:0 and 2:1 D1D2: AVBS1-2 or 3:0 and 3:1 D1D2:AVBS1-3 complexes. However, the discrepancy between the determined and expected molecular masses of the complexes, as well as the slopes observed for the molecular mass argued that the peak corresponded to 2:0 and 2:1, or 3:0 and 3:1 complexes in equilibrium (Figs. 2 d-f). In line with this, quantitative SDS-PAGE analysis of the peak fractions indicated a molar ratio comprised between 2:0-2:1 and 3:0-3:1 for D1D2:AVBS1-2 and D1D2:AVBS1-3, respectively, consistent with a mixture of inter-exchanging complexes present in the corresponding peaks (Fig. S2).

These results suggest that binding of IpaA VBS1-3 to vinculin triggers conformational changes leading to the formation of vinculin trimers.

### IpaA promotes major conformational changes in the vinculin D1 D2 domains

To further investigate mechanism responsible for vinculin oligomerization, we performed binding assays with vinculin derivatives immobilized onto a solid phase to restrict conformational changes. By constraining conformation of vinculin derivatives, we expected to prevent the formation of higher order oligomers observed in solution, while enabling binding of IpaA VBSs to initial sites on the vinculin derivative conformers. These assays indicated that AVBS1-3 and AVBS1-2 bound to vinculin with a similar affinity as estimated in Fig. S3a by their EC50 (95% confidence interval) of 6.1 (4.2-9.0) and 3.7 (1.7-8.1) nM, respectively. Strikingly, a large difference was observed in the binding plateau, indicating that vinculin presented more binding sites for AVBS1-3 than for AVBS1-2 (Fig. S3a). D1D2 presented more binding sites than the D1 domain only, suggesting the presence of additional sites on the D2 domain (Fig. S3b). Consistently, BN-PAGE showed the formation of 1:1, as well as a 1:2 D1D2:AVBS1-3 complexes, observed with increasing AVBS1-3 molar ratio (Fig. S3c). In contrast, single 1:1 complexes were observed for D1:AVBS1-3, D1:AVBS1-2 or D1D2-AVBS1-2 (Figs. S3c-h), indicating that IpaA VBS3 was required to reveal additional sites on the D2 domain. These results suggested that as for immobilization on solid phase, the presence of Coomassie brilliant blue in BN-PAGE interfered with higher order vinculin oligomerization while enabling binding to the vinculin derivative monomer. Together, these results suggested that the formation of vinculin oligomers triggered by AVBS1-3 required the IpaA VBS3 dependent exposure of binding sites on D2. These findings were unexpected, since vinculin activating ligands have been described to bind to a single site on the D1 domain of vinculin.

To map interactions between AVBS1-2 and AVBS1-3 with D1D2, complexes were crosslinked, subjected to proteolysis and analyzed using Liquid Chromatography coupled to Mass Spectrometry (LC-MS) (Star Methods). Intermolecular links were identified from the characterization of cross-linked peptides, and along with identified intramolecular links, used to produce structural models (Suppl. Tables 1-3 and Figs. S4a, b; Star Methods). The D1:AVBS1-2 complex showed links consistent with a “canonical” conformer expected from established structures (Izard, Tran Van Nhieu et al. 2006, Tran Van Nhieu and Izard 2007) (Fig. S4c). Similar links were identified for the D1D2:AVBS1-2 complex, with a majority of links observed with the D1 domain (Fig. S4d). For both complexes, the structure shows interactions between IpaA VBS1 and VBS2 with the D1 first and second bundles, respectively, leading to helical bundle reorganization of D1 associated with vinculin activation (Izard, Evans et al. 2004; Figs. S4c, d). For the D1D2:AVBS1-3 complex, MS-based structural modeling reveals two major conformers accounting for the majority of links. In a first “closed” conformer, IpaA VBS1 and VBS2 interact with the D1 bundles in a similar manner as for AVBS1-2, where the relative positioning of D1 and D2 is globally conserved compared to apo D1D2 or the D1D2:AVBS1-2 complex (Fig. 3a and Figs. S4d, e). In this “closed” conformer, IpaA VBS3 interacts with an interface formed by the H5 (residues 128-149) and H8 (residues 222-250) helices in the second bundle of D1, and the H13 (residues 373-397) helix in the second bundle of D2 (Figs. 3a, b). The second “open” conformer, however, shows a major re-orientation of D1 and D2 subdomains with their major axis forming an angle value of ca 82° compared to the 25° observed in the native vinculin structure or the first conformer, with IpaA VBS3 docking sidewise through extensive interaction with the H5 (residues 128-149) and H8 (residues 222-250) helices of D1 (Figs. 3c, d, light blue helices). Since this latter conformer was observed for AVBS1-3 but not for AVBS1-2, we posited that it was involved in the formation of higher order D1D2 complexes and homotrimer. To test this, we engineered mutations substituting residue Q68 in the first D1 bundle and A396 in the second D2 bundle for cysteine residues, expected to prevent the formation of the open conformer upon disulfide bridge formation (Fig. 3e) by preventing major conformational shifts of the D1 and D2 domains. In control experiments, disulfide bridge formation was detected in D1D2 and full-length HV containing the Q68C and A396C mutations, expected to act as a clamp preventing the major conformational changes induced by AVBS1-3 (Figs. S5a-c).

**Figure 3.**
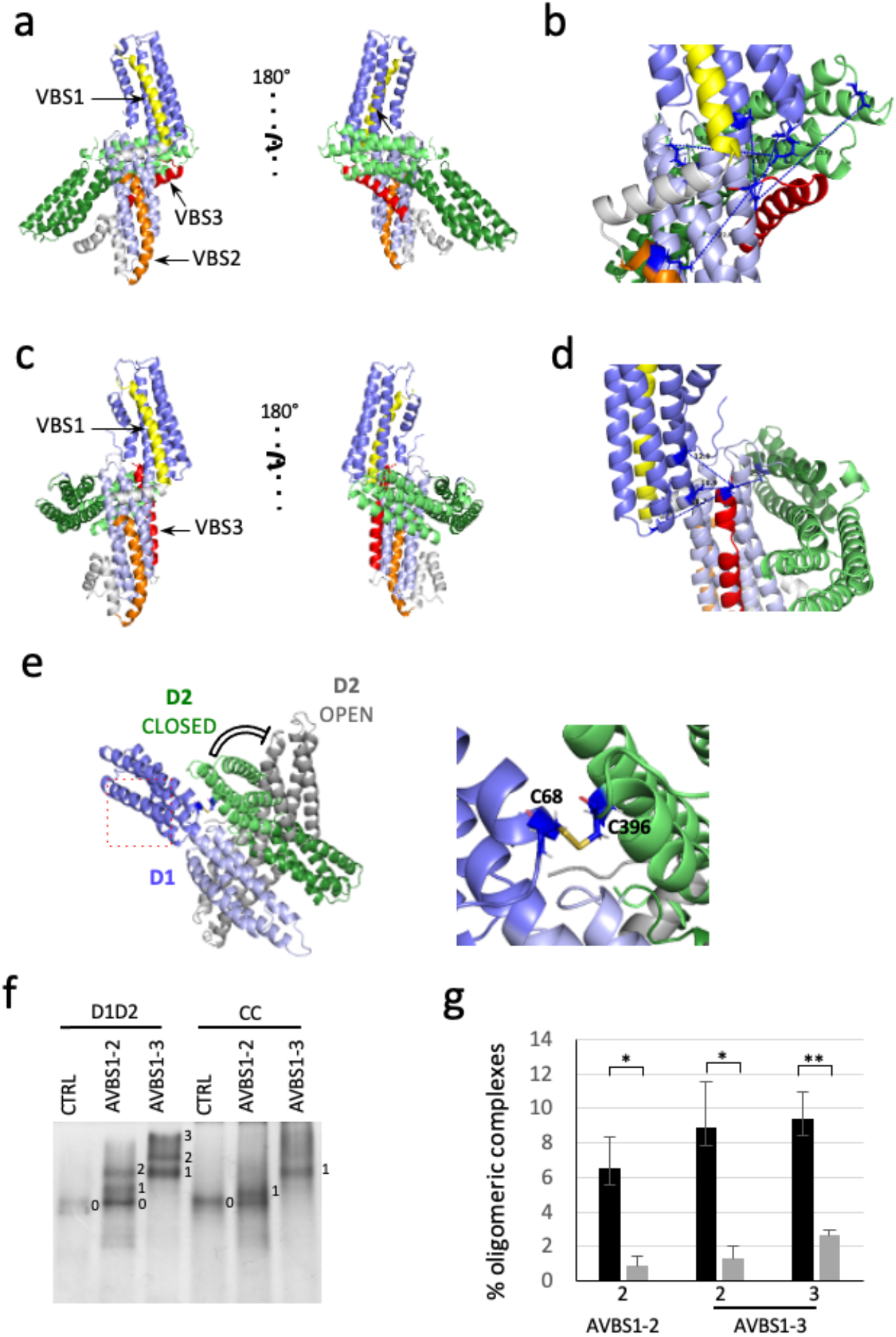
IpaA promotes major conformational changes in the vinculin D1 D2 domains. **a-e**) Structural models of D1D2-IpaA VBS1-3. Pale blue: D1 first bundle. Dark blue: D1 second bundle. Pale green: D2 first bundle. Dark green: D2 second bundle. Yellow: IpaA VBS1. Orange: IpaA VBS2. Red: IpaA VBS3. **a, b**, “closed” conformer; **c, d**, “open” conformer. **h, j**, higher magnification of the IpaA VBS3-D1D2 interaction in (*b*) and (*d*) showing the identified cross-linked distance between residues in Å. IpaA VBS1-3 were docked on the surface of Vinculin D1D2 and verified using MS crosslink constraints. TX-MS protocol (Hauri, Khakzad et al. 2019) in combination with MS constraints was used to unify and adjust the final model, which justifies over 100 cross-links. e, Structural model of cysteine-clamped vinculin. Green: D2 in the closed conformer. Grey: D2 domain in the open conformer. Black: C68-C396 cysteine clamp preventing the switch from closed to open conformers. Right panel: enlarged view of the cystein clamp shown in the inset in the left panel. **f**, native gel analysis of vinculin D1D2 and IpaA derivatives. D1D2 or the double cysteine mutant D1D2 (CC) were incubated with the indicated IpaA derivatives and analyzed by native PAGE followed by Coomassie staining. The numbers next to the band indicate upper-shifted bands of D1D2:AVBS1-2 or D1D2:AVBS1-3 complexes. Note the absence of higher order complexes for the CC mutant. **g**, the band integrated intensity corresponding to the indicated shifted bands were quantified using ImageJ. Values are expressed as the percent of total protein amounts in the corresponding sample. Solid bars: D1D2. Grey bars: CC.

As shown in Fig. 3f, two and three upper-shifted D1D2 bands were visualized by clear native PAGE upon incubation with AVBS1-2 and AVBS1-3, respectively. Quantitative 2^nd^ dimension SDS-PAGE analysis of the upper-shifted bands 2 and 3 upon incubation with AVBS1-3 indicated a D1D2:AVBS1-3 molar ratio superior to 3, suggesting these likely corresponded to the 2:0-2:1 and 3:0-3:1 higher order D1D2 complexes observed in the SEC-MALS analysis (Figs. 2, S2 and S5e), although with a different representativity perhaps linked to electrophoretic conditions. The cysteine clamp Q68C A396C (CC) in D1D2 did not prevent the exposure of additional sites on D2 or 1:1 complex formation induced by AVBS1-2 or AVBS1-3. However, CC prevented the formation of higher order complexes for D1D2 as well as for full-length vinculin (Figs. 3f, and S5f, HV-CC). We coined “supra-activation” the mode of vinculin activation induced by AVBS1-3 involving major conformational changes in the vinculin head to distinguish it from the canonical activation associated with the dissociation of vinculin head-tail domains.

### IpaA mediates actin bundling and vinculin-talin co-clusters at a distance from activation site

To further characterize the role of vinculin “supra-activation”, we performed actin cosedimentation assays. As expected, the cysteine-clamp had little effects on vinculin canonical activation, since the majority of cysteine-clamped full-length vinculin (HV-CC) associated with actin filaments upon incubation with AVBS1-2 or AVBS1-3 (Figs. S6a, b). We then tested the ability of vinculin oligomers to promote actin bundling by performing low-speed sedimentation assays (Star Methods). As shown in Fig. 4, vinculin alone did not promote actin bundling (Figs. 4a, b, HV). Upon incubation with AVBS1-3, up to 30% of the total actin pool sedimented, consistent with AVBS1-3-mediated vinculin actin bundling. This actin bundling activity was associated with the low-speed co-sedimentation of vinculin with actin (Fig. 4a). In contrast, no such actin bundling activity was observed for HV-CC even upon incubation with AVBS1-3 (Figs. 4a, b). Together, these results suggest that vinculin oligomers triggered by IpaA-mediated vinculin supra-activation bundle actin filaments.

**Figure 4.**
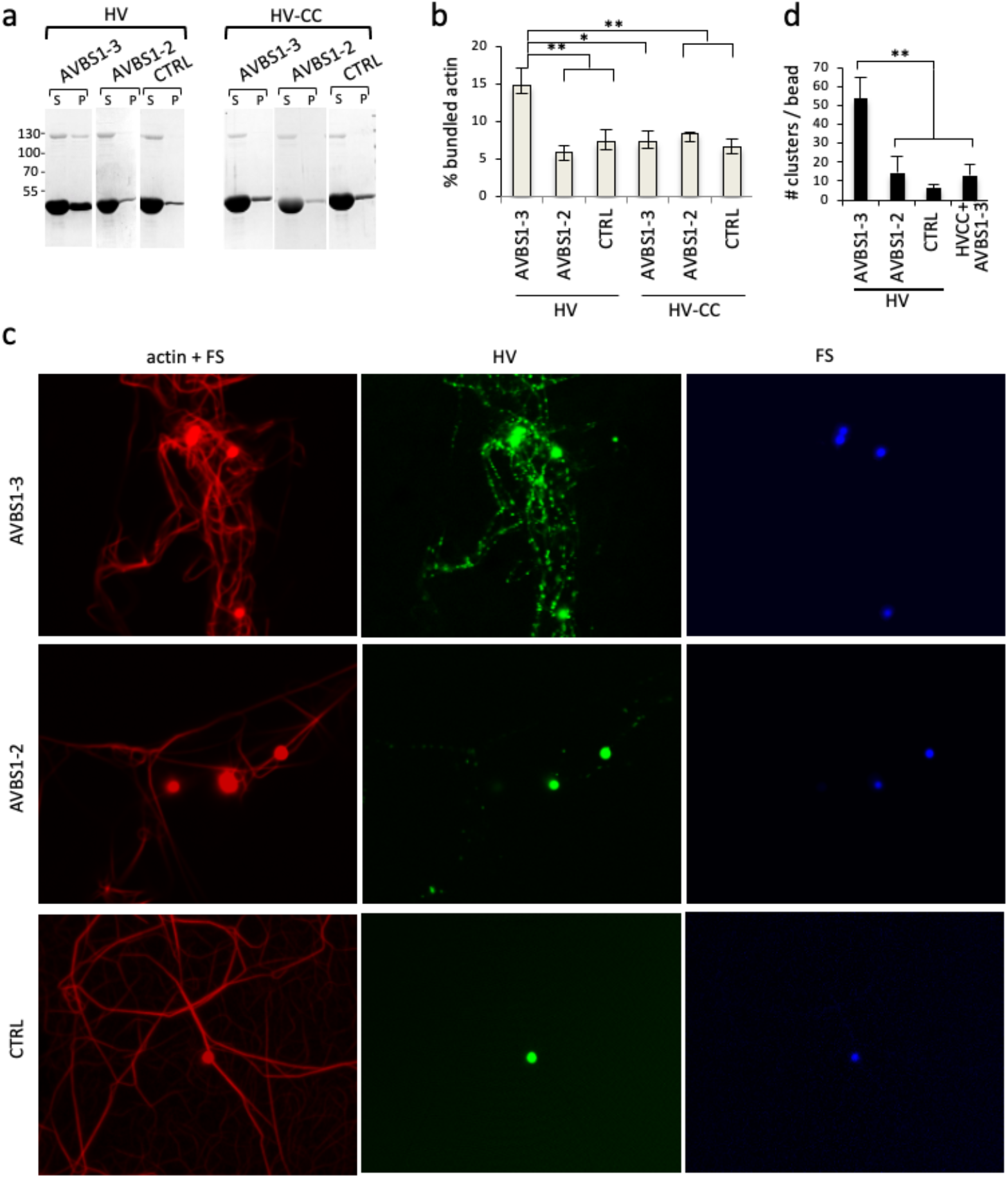
IpaA-induced vinculin oligomers promote actin bundling at a distance from activation sites. **a, b,** actin sedimentation assays. HV: full-length human vinculin; HV-CC: cysteine-clamp derivative. Actin was allowed to polymerize at a final concentration of 20-30 μM in the presence of the indicated proteins. Samples were centrifuged at 14,000 g for 10 min to pellet actin bundles, respectively. **a**, representative SDS-PAGE analysis using a 10% polyacrylamide gel followed by Coomassie staining. S: supernatant; P: pellet. **b**, the band integrated intensity corresponding to actin was quantified using ImageJ. Values are expressed as the percent of actin in the pellet fraction relative to the total actin amounts in the supernatant and pellet fractions normalized to control. **b**, percent of bundled actin; HV (n = 9, N = 4); HV-CC (n = 6, N = 3). **c**, Representative micrographs of fluorescently labeled actin polymerized in the presence of Bodipy-vinculin and coated beads. Beads were coated with AVBS1-3, AVBS1-2 or GST (CTRL). red: actin; green: Bodipy-vinculin; blue: beads. Numerous vinculin clusters are observed at a distance from AVBS1-3 coated-beads. **d**, the numbers of vinculin clusters per bead ± SEM are shown for the indicated samples (N= 3, HV+AVBS1-3: 1292; N= 2, HV+AVBS1-2: 120; N=2, HV+GST (CTRL): 56; N=2, HVCC+AVBS1-3: 80). Scale bar = 5 μm. Mann and Whitney. *: p < 0.05; **: p < 0.01.

Next, we asked how vinculin oligomers promoted actin bundling relative to the site of AVBS1-3-mediated supra-activation. Indeed, vinculin homo-oligomers are not expected to remain bound to AVBS1-3 but to diffuse away from activation sites. To test this, we designed a solid-phase assay where GST-AVBS1-3 was coated on 1 μM-diameter fluorescent beads (Star Methods). Control experiments indicated that GST-AVBS1-3 showed little desorption from beads up to 2 hours following coating (Figs. S6c, d). HV was fluorescently labeled and incubated with GST-AVBS1-3-coated beads in actin polymerization assays (Star Methods). As shown in Fig. 4c, vinculin clusters were clearly detected in association with actin bundles, away from AVBS1-3-coated beads. As expected, such vinculin clusters were observed to a much lesser extent with beads coated AVBS1-2, control GST, and the HV-CC cysteine clamp construct (Figs. 4c, d). Also, consistent with low-speed actin sedimentation results, actin bundling was prominent upon incubation with AVBS1-3-coated beads, relative to AVBS1-2- and GST-coated beads and was not observed for HV-CC (Figs. 4c, d). These results are consistent with the formation of vinculin homo-oligomers induced by AVBS1-3-mediated supra-activation, bundling actin filaments away from the activation sites.

### Talin VBSs promote higher order clustering of IpaA-induced vinculin oligomers

During canonical activation, vinculin simultaneously binds to talin and actin filaments through its N-terminal and C-terminal domain. We then used fluorescent vinculin clustering assays to ask whether in addition to actin bundling, AVBS1-3-mediated vinculin oligomers could interact with talin. In the design of these experiments, we aimed to mimic the multiplicity of VBSs present per talin molecule expected to play an additional scaffolding role in vinculin cluster formation at FAs. For this purpose, we coated 100 nm-beads with the vinculin binding H1-H4 helices from the R1 talin bundle at a calculated density of one H1H4 molecule / 86 nm^2^ and co-incubated these beads along with AVBS1-3-beads and vinculin in actin polymerization assays. As shown in Fig. 5a, Tln-beads co-localized with vinculin clusters induced by AVBS1-3-coated beads consistent with binding of vinculin oligomers to talin. As expected, very few clusters were observed for AVBS1-2-coated beads (Fig. 5a). In addition, the integrated density of vinculin clusters showed striking difference between AVBS1-2 and AVBS1-3, with AVBS1-3-induced clusters being on average 9-times brighter than AVBS1-2-induced clusters (Figs. 5a, left panels and 5b). These marked differences suggested additional clustering levels mediated by multivalent talin beads. To confirm this, we quantified the density of Tln-beads per vinculin clusters based on their integrated fluorescence intensity. As shown in Fig. 5c, talin beads showed a recruitment that was 2.3-fold higher at vinculin clusters induced by AVBS1-3 relative to AVBS1-2, suggestive of higher order clustering.

**Figure 5.**
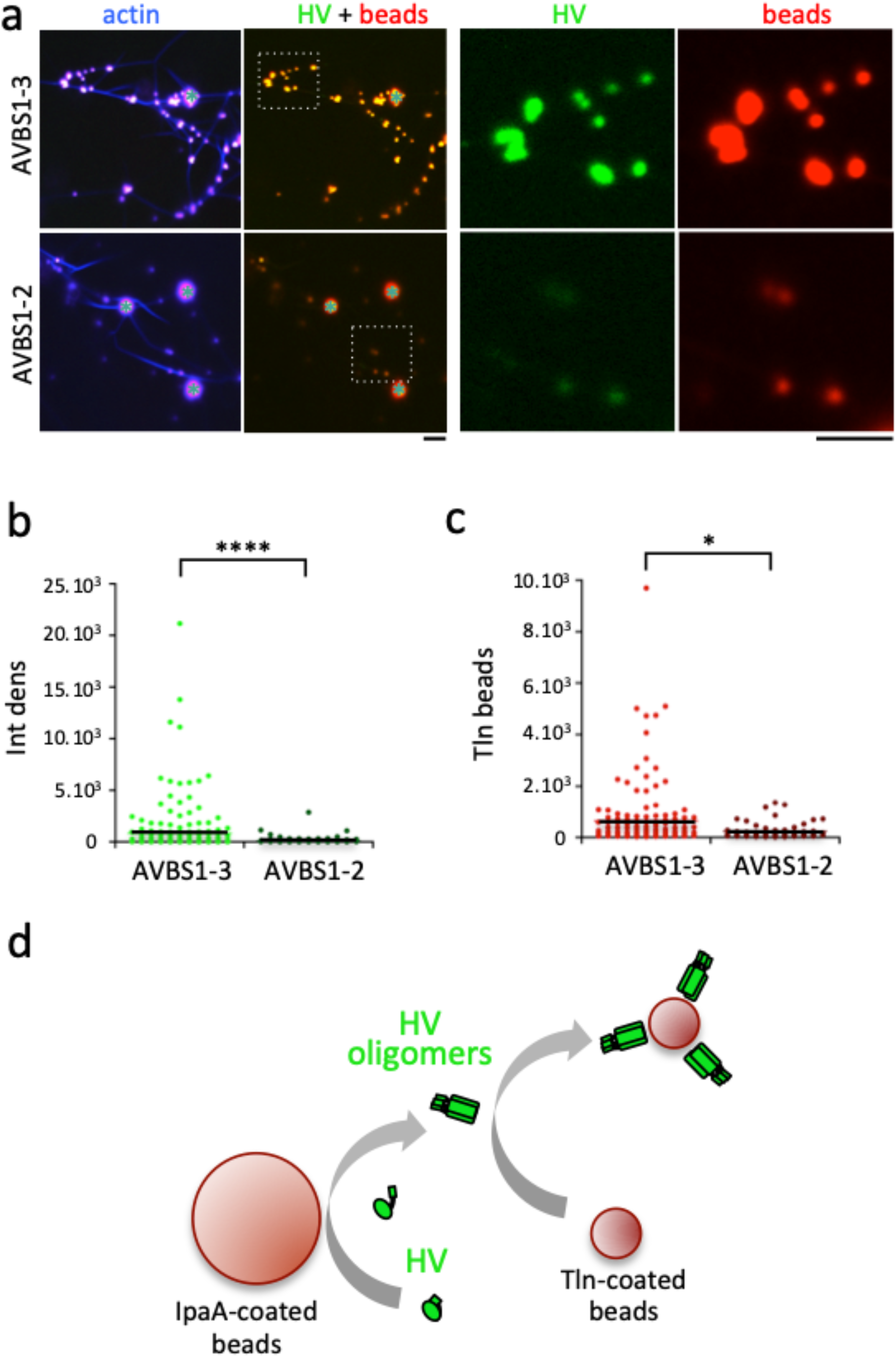
Vinculin oligomers are clustered by talin VBSs. Fluorescent actin was allowed to polymerize in the presence of vinculin, 1 μm-diameter red fluorescent beads coated with the indicated sample and 100 nm-diameter red fluorescent beads coated with talin H1-H4. **a**, representative micrographs. Blue: actin. Green: vinculin; red: 1 μm- and 100 nm-red fluorescent coated beads. The cyan stars indicate GST-AVBS1-2 / GST-AVBS1-3 –coated 1 μm beads. Right panels are higher magnification of the insets shown in the left panels. Scale bar = 1 μm. **b**, normalized integrated density of vinculin clusters. HV+AVBS1-3: 204 clusters, N =2; HV+AVBS1-2: 142 clusters, N =2. **c**, normalized integrated density talin H1-H4 beads. HV+AVBS1-3, 148 clusters, N =2; HV+AVBS1-2: 64 clusters, N =2. Mann and Whitney. *: p = 0.023; ****. p = 0.00018. **d**, model for vinculin oligomers clustering by talin VBSs. Bead-immobilized AVBS1-3 catalyzes the formation of vinculin oligomers, that diffuse away from activation site. Vinculin oligomers binding to Tln-coated beads results in higher order cluster formation.

Together, the results indicate that as opposed to canonical activation, vinculin supra-activation leads to the formation of homo-oligomers mediating actin bundling and binding to talin, a property promoting another levels of clustering by multivalent VBSs. Following diffusion, vinculin oligomers show persistent F-actin binding and bundling activity at a distance from the activation site.

### Vinculin supra-activation promotes actin bundling and adhesion expansion

We next tested the effects of vinculin supra-activation on FA formation by introducing the cysteine clamp in full length vinculin fused to mCherry (CC-HV) and analyzed its effects following transfection in MEF vinculin-null cells. As shown in Fig. 6a, vinculin led to the formation of larger and more numerous talin-containing FAs than mock-transfected vinculin null cells, consistent with residual vinculin activation (Figs. 6a, b). In contrast, CC-HV-expressing cells formed significantly fewer and smaller FAs than cells transfected with vinculin (Figs. 6a-e). A more detailed analysis indicated that the average width of adhesions formed by CC-HV was remarkably conserved with an average of 0.96 ± 0.27 (SD) μm (Figs. 6f, h, i). This was in sharp contrast with FAs formed by wildtype vinculin showing a larger dispersion in average width (1.31 ± 0.57 (SD) μm) and reaching up to several microns (Figs. 6g-i). In addition, stress fibers and thick actin bundles connecting FAs were observed for HV, but not CC-HV expressing cells (Figs. 6a, f, g). These results suggest that vinculin supra-activation impaired in CC-HV, is involved in the growth of adhesion structures and actin bundling during FA maturation.

**Figure 6.**
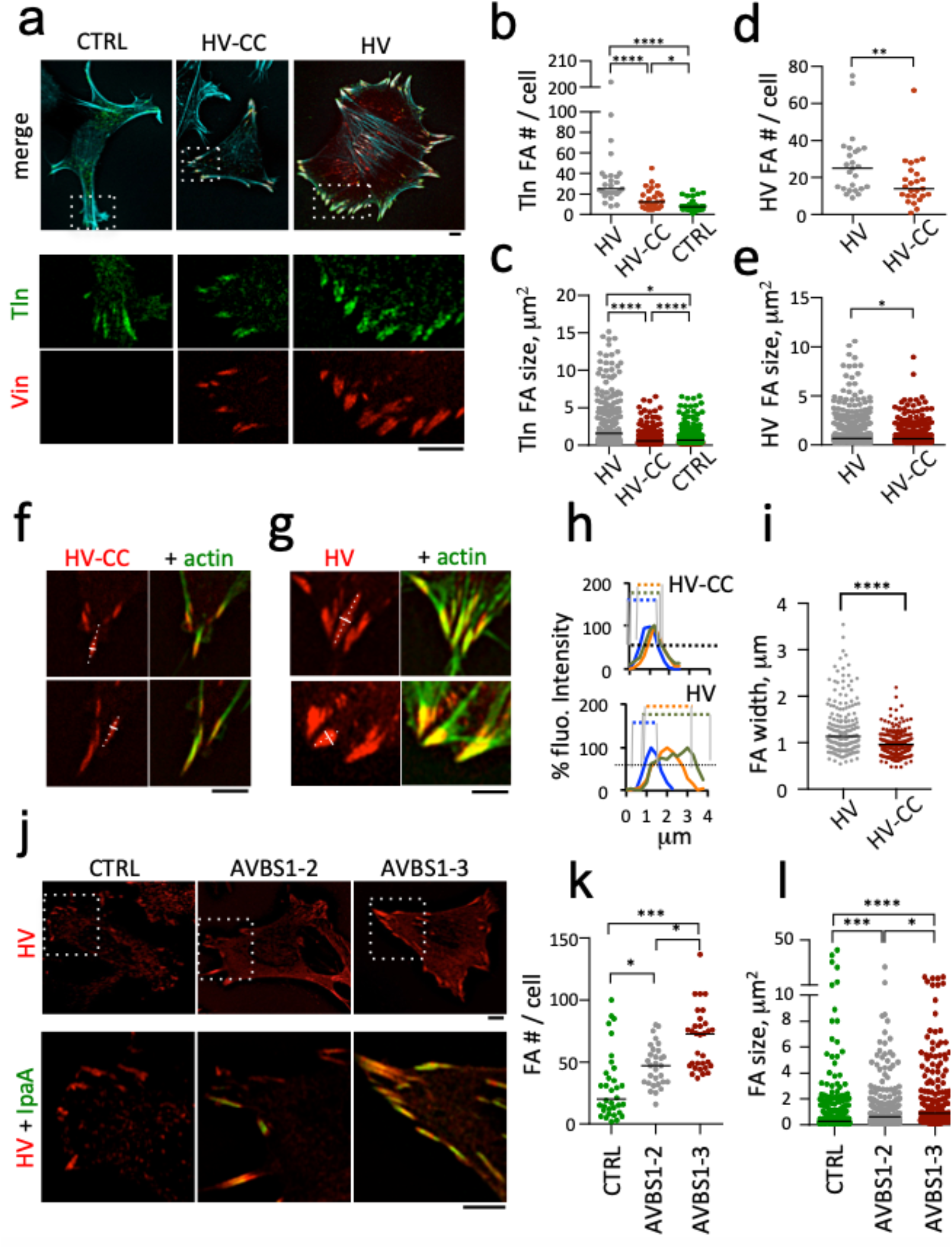
vinculin supra-activation promotes the merging of adhesions clusters. Cells were transfected, fixed and processed for fluorescence microscopy analysis of FAs. **a, f, g, j**, representative fluorescence micrographs; red: Vinculin-mCherry; green: GFP-talin (**a**) or F-actin (**f, g**); cyan: F-actin (**a**). Scale bar = 5 μm. **a**-**i**, MEF vinculin null cells; transfection with GFP-talin (CTRL); talin co-transfection with full-length HV-mCherry (HV) or HV Q68C A396C-mCherry (CC-HV). **j-l**, C2.7 cells; transfection with vinculin (CTRL); vinculin co-transfection with GFP-AVBS1-2 or GFP-AVBS1-3. **b-e, k, l**: the FA number per cell and size were determined using a semi-automatic detection program (Star Methods). Bar: median size. FAs analyzed for: **b**, **c**, GFP-talin; **d**, **e, h**, **i**, **k, l,** HV or CC-HV. **b-e**, CTRL: n=28, N=3; HV: n = 25, N = 3; CC-HV: n = 25, N = 3. **h**, representative plot profiles from linescans (solid white lines) orthogonal to the main FA axis (dashed white lines) in (**f**) and (**g**). **i**, FA width determined as the full width half-maximum by linear interpolation from plot profiles in (**h**); HV: 181 FAs, 6 cells, N = 2. CC-HV: 101 FAs, 14 cells, N = 3. Mann-Whitney test with Bonferroni multiple comparison correction. *: p < 0.05; **: p < 0.01; ***: p < 0.005; ****: p < 0.001. **j-l**, n > 30 cells, N = 3. Dunn’s multiple comparisons test. *: p < 0.05; ***: p < 0.005.

To clarify the role of IpaA-mediated vinculin supra-activation in FA dynamics, we analyzed the effects of AVBS1-2 and AVBS1-3 expression in C2.7 cells, a vinculin-expressing myoblastic cell line, which form prominent FAs well suited for dynamic TIRF (total internal reflection fluorescence) microscopy analysis. As shown in Figs. 6j-l and S6a, b, cells transfected with GFP-AVBS1-2 formed more numerous and larger peripheral FAs as well as actin-rich ruffles compared to control cells. GFP-AVBS1-3 transfected cells formed even larger and more numerous FAs, but with significantly less actin ruffles than GFP-AVBS1-2 transfected cells (Figs. 6j-l and S7a, b). Strikingly, GFP-AVBS1-3-induced FAs were extremely stable, with a median duration of at least 84 min, while GFP-AVBS1-2-transfected and control cells showed FAs with a comparable median duration of less than 25 min (Figs. S7c, d; Suppl. movie 1). This increased FA stability in GFP-AVBS1-3 transfectants was predominantly due to decreased rates of FA disassembly with a 2-fold decrease in median instant rates relative to control cells (Figs. S7e, f; Suppl. movie 1). These results indicate that AVBS1-3-induced vinculin supra-activation promotes the expansion and increased stability of FAs.

### IpaA-induced focal adhesions form independent of mechanotransduction

The stability of IpaA-induced FAs suggests that their formation may be less dependent on mechanotransduction. To test this, we analyzed the effects of the acto-myosin relaxing Rho-kinase inhibitor Y27632. Strikingly, vinculin-labeled FAs induced by GFP-AVBS1-3 resisted the action of Y27632, with a five- and four-times slower median rate of FA disassembly relative to control cells and GFP-AVBS1-2 transfectants, respectively (Figs. 7a-c Suppl. movie 2). Large FAs were even observed to form in GFP-AVBS1-3 transfectants following addition of the inhibitor (Figs. 7a-c), a process that was not observed for other samples, including cells transfected with GFP fused to the vinculin D1 domain (vD1) reported to delay talin refolding following stretching (del Rio, Perez-Jimenez et al. 2009, Margadant, Chew et al. 2011, Carisey, Tsang et al. 2013) (Figs. 7a-c; Suppl. movie 2). GFP-AVBS1-3 also delayed the Y27632-induced removal of the late adhesion marker VASP (Fig. S8; Suppl. movie 3).

**Figure 7.**
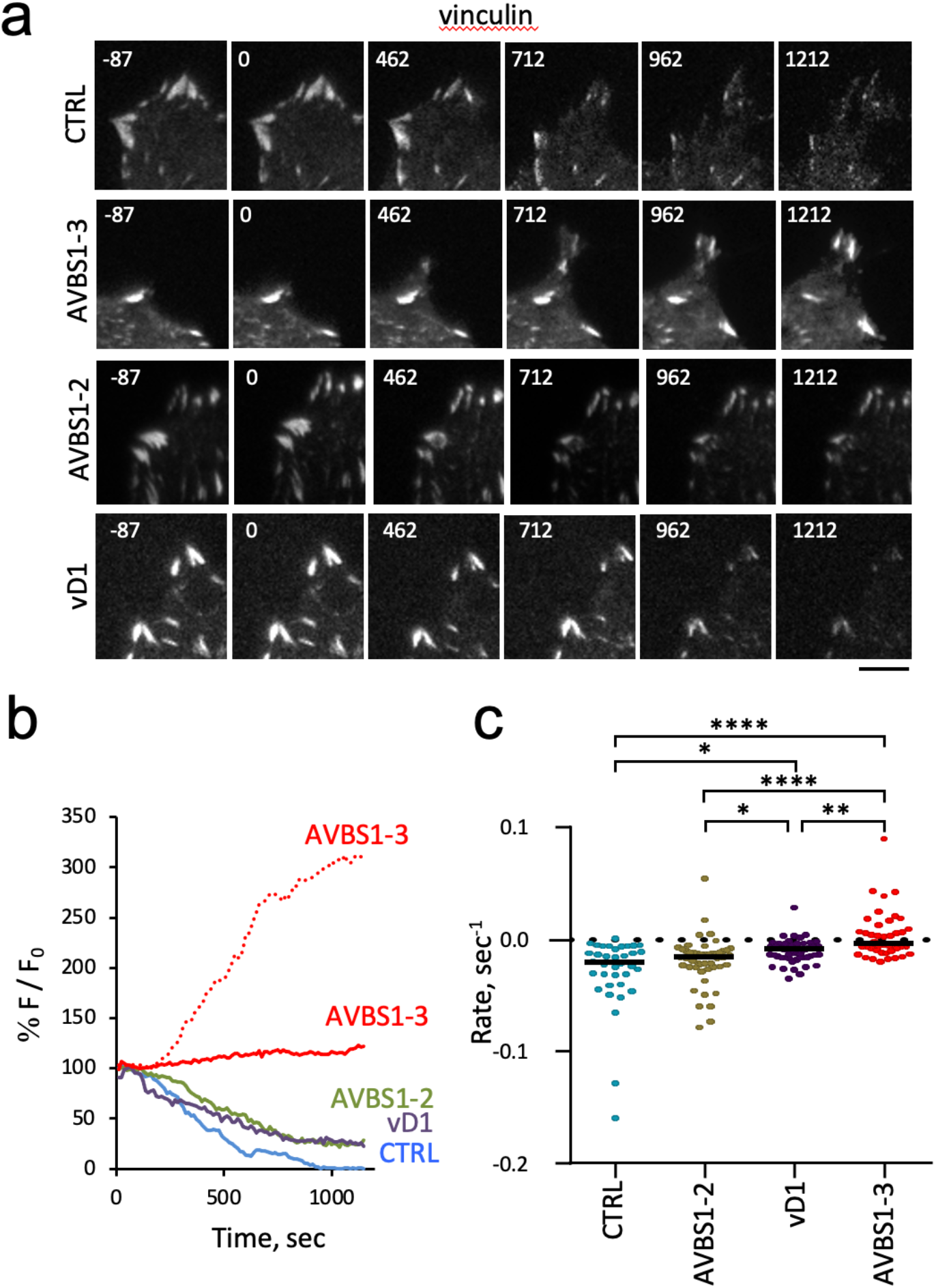
IpaA-mediated vinculin supra-activation stimulates cell adhesion independent of mechanotransduction. TIRF microscopy of C2.7 cells transfected with mCherry-vinculin alone (CTRL) or co-transfected with GFP-AVBS1-2 (AVBS1-2), GFP-vD1 (vD1), or GFP-AVBS1-3 (AVBS1-3). Adhesion kinetic parameters were determined from time-lapse acquisitions following cell treatment with 100 μM Y-27632. **a,** representative time series acquisitions. Numbers indicated the elapsed time in seconds, with the inhibitor added at t = 0. Scale bar = 5 μm. **b**, % F/F_0_: average fluorescence intensity of adhesions expressed as a percent of initial fluorescence. Solid lines: representative traces corresponding to single adhesions for the indicated samples in the corresponding color. The dashed redline illustrates FA assembly in GFP-AVBS1-3 transfected cell, seldom observed with the other samples. **c**, initial rates of adhesion assembly / disassembly inferred from linear fits. Number of adhesions analyzed: **c**, N = 5. CTRL: 84; vD1: 75; AVBS1-2: 140; AVBS1-3: 97. Dunn’s multiple comparisons test. *: p < 0.05; ****: p < 0.001.

These findings are consistent with our in vitro results showing the catalysis by IpaA of vinculin oligomers with persistent activity. The resistance of VASP to the action of Y27632 in IpaA-induced FAs suggests that vinculin oligomers contribute to the scaffolding of this late FA marker.

## Discussion

*Shigella* invades host cells through a triggering mode implicating discrete number of contacts between the T3SS and host cells (Valencia-Gallardo, Carayol et al. 2015). How IpaA promotes bacterial attachment to the cell surface by reinforcing cytoskeletal tethering to limited bacterial contact sites has been an open question. Here, we show that IpaA induces the supra-activation of vinculin associated with unveiling of binding sites on the D2 subdomain and major conformational changes of the vinculin head. Vinculin supra-activation leads to the formation of vinculin homooligomers that bundle actin filaments and bind to talin. Strikingly, IpaA-induced vinculin oligomers diffuse away from activation sites, a property associated with the expansion of adhesion structures at bacterial contact sites during *Shigella* invasion. Our studies also suggest that vinculin supra-activation is also involved in the maturation of cell adhesions, independent of bacterial invasion: i) a cysteine-clamp inhibiting vinculin supra-activation, but not canonical activation prevents the formation of mature adhesions; ii) IpaA VBS1-3 that mediates vinculin supra-activation accelerates the speed of cell adhesion but at a steady state, the strength of cell adhesion does not differ from that of control cells. These results suggest that IpaA VBS1-3 mediates vinculin supra-activation through the unique organization and joint action of its three VBSs, but that canonical activation also leads to supra-activation when combined with mechanotransduction. Consistent with this, we also found that during *Shigella* invasion, vinculin supra-activation triggered by IpaA VBS1-3 is required for full adhesion formation at bacterial contact sites, canonical activation mediated by a single IpaA VBS is sufficient for FA formation at basal membranes distal from bacteria.

Interestingly, molecular dynamics simulations suggest that stretching of the vinculin during mechanotransduction also results in the exposure of the vinculin head D2 subdomain (Kluger, Braun et al. 2020). It is therefore tempting to speculate that mechanotransduction also leads to vinculin homo-oligomerization that we observed for IpaA through intermolecular interactions between the D1-D2 domains. Whether vinculin stretching alone, or an additional interaction between a VBS and D2 in a manner similar to IpaA VBS3 is required during mechanotransduction is not known. Of interest, IpaA VBS3 shares homology with the talin VBS corresponding to helix 46 (Valencia-Gallardo, Bou-Nader et al. 2019). This suggests that talin H46 helix could play such a role during vinculin supra-activation.

There are important differences, however, that one can expect from vinculin supra-activation depending on IpaA or mechanotransduction. These differences could account for the extreme stability of FAs in IpaA VBS1-3-compared to IpaA VBS1-2-transfectants during cell treatment with actin relaxing drugs. Indeed, IpaA acts as a catalyst leading to the persistent production of vinculin oligomers independent of the stretching of vinculin by cytoskeletal forces. During vinculin canonical activation, however, oligomers will form as a function of the stretching force and disassemble when the force is released.

Vinculin oligomers are sufficiently biochemically stable to be observed in our SEC-MALS, native gel or fluorescence microscopy analysis, suggesting the stabilization of conformers via inter-protomer interactions. Because of this property, vinculin supra-activation may correspond to a switch defining a threshold during the formation of adhesion structures. Indeed, because of their relative stability, their ability to simultaneously bundle actin filaments, bind to talin and to diffuse away from activation sites, vinculin oligomers following supra-activation are expected to promote the expansion and strengthening of adhesion structures through diffusion and capture by talin VBSs. Consistently, our studies show that supra-activation-deficient cysteine clamped vinculin only support the formation of small adhesion structures (< 1 μm width) with limited actin bundling, while supra-activation proficient vinculin promotes the formation of large FAs and actin bundles. Our talin-bead scaffolding assays indicate that vinculin cluster formation is further driven by the multiplicity of talin VBSs (Fig. S7, grey arrows). Because of the multiplicity of VBSs on talin molecules, co-clustering between vinculin oligomers and talin is likely to play a role in adhesion expansion (Fig. S7). Vinculin- has been shown to immobilize and bundle actin filaments from Arp2/3 branched networks (Boujemaa-Paterski, Martins et al. 2020). In this case, however, bundling was shown to occur at activation sites and proposed to occur through talin-vinculin scaffolds (Boujemaa-Paterski, Martins et al. 2020). This activity is in contrast with diffusible bundling activity associated with vinculin oligomers that we describe here, which may promote clustering at a distance from activation sites. Clustering of adhesions at different scales is believed to play a major role in adhesive processes through the regulation of functional units (Mege and Ishiyama 2017). At the molecular levels, integrin clustering in adhesions is not fully understood; it may be induced by ligand-binding and integrin homo-oligomerization via integrin trans-membrane domains (Karimi, O’Connor et al. 2018). Nanoclusters consisting of hundreds of integrin molecules were proposed to correspond to elementary units that merge to form nascent adhesions (Changede and Sheetz 2017). Vinculin oligomers following supra-activation could provide an additional basis for the expansion of adhesions through the bundling of actin filaments and scaffolding of cytosketal linkers in response to increasing acto-myosin pulling force (Fig. S9). As opposed to physiological substrates, single bacteria cannot sustain the range of counterforces associated with the strengthening of adhesion structures during mechanotransduction and integrin-mediated adhesion to the substrate. Actin bundling and scaffolding triggered by IpaA-induced vinculin oligomers may *Shigella* with means to strengthen its adhesion during bacterial invasion independent of mechanotransduction (Fig. S9a). Importantly, the view of IpaA catalyzing the formation of vinculin activated vinculin oligomers may explain its potency and how a limited number of this injected type III effector can have major impact on the global cell adhesion properties, distal from bacterial invasion sites (Fig. S9b). Understanding how vinculin supra-activation and oligomer formation regulate the composition and properties of adhesions at *Shigella* invasion sites will likely have important implications for cell adhesion and will be the focus of future investigation.

### Limitations of the study

We describe a novel mode of vinculin supra-activation induced by the Shigella type III effector IpaA. Whether vinculin supra-activation also occurs during cell adhesion during the maturation of adhesion structures will required further investigation.

## ACKNOWLEDGEMENTS

The authors thank Gauthier Mercante for technical help, Philippe Mailly for help with image analysis and René-Marc Mège for insightful discussions and reading of the manuscript. This work was supported by grants from Inserm, CNRS and Collège de France to the CIRB, as well as grant from the PSL Idex project “Shigaforce”. DIAS and BCC are recipients of a PhD fellowship from a CONACYT scholarship. CV-G and DIAS also received support from the Memolife Labex. HK and LM were supported by Swiss National Science Foundation (grant no. SNF P2ZHP3_191289) and the Knut and Alice Wallenberg Foundation (grant no. KAW 2016.0023), respectively. This work has also benefited from the facilities and expertise of the Macromolecular Interaction Platforms of I2BC.

## AUTHOR CONTRIBUTIONS

CV-G and DIAS conceived and performed most of the experimental works, data analysis and wrote the manuscript. BCC, SE and BM analyzed TIRF experiments. CV and CB-N performed the SEC-MALS analysis. BCC and YZ performed native gel analysis of vinculin complexes. CM and JCR designed and performed the LC-MS analysis. DBL analyzed the cross-linked mass spectrometry data. HK and LM generated structural models. CL provided help for the actin co-sedimentation assays. GTVN designed the project and wrote the manuscript.

