## Supplementary Materials for "*Shigella* ipaA mediates actin bundling through diffusible vinculin oligomers with activation imprint"

**SUPPLEMENTARY INFORMATION**

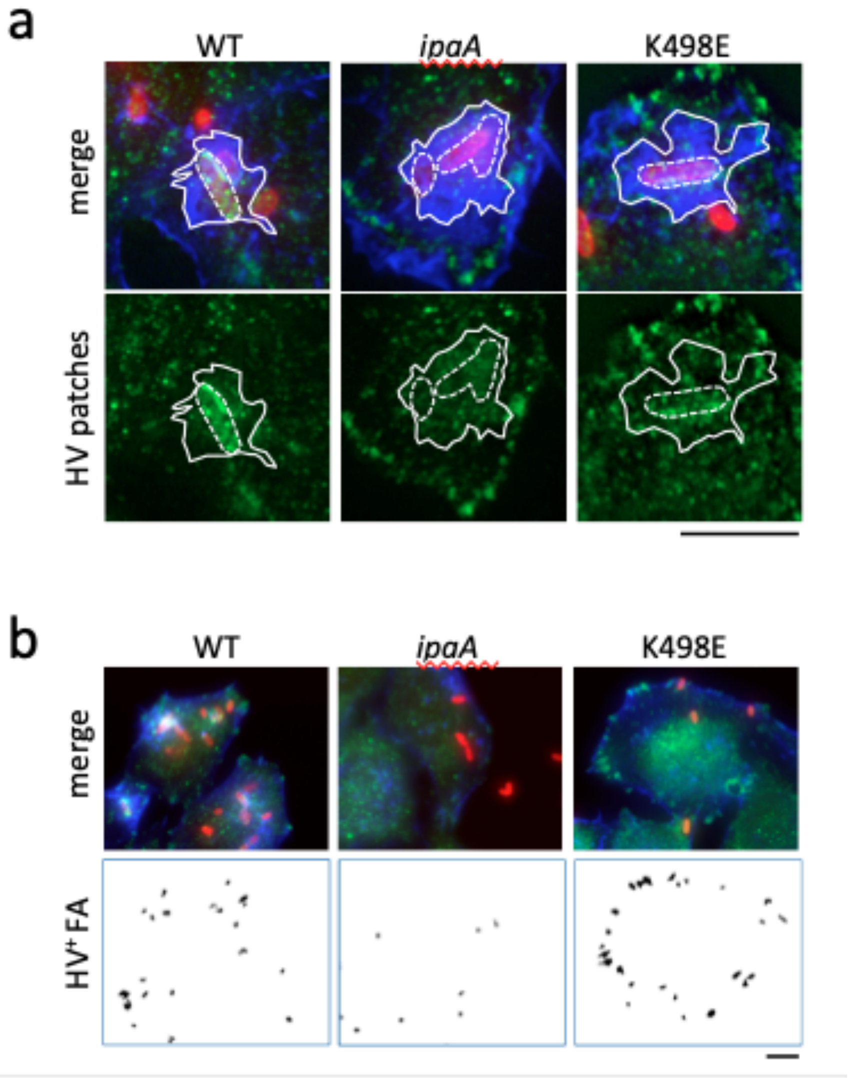

**Supplementary Fig. 1. Quantification of IpaA-dependent vinculin recruitment during *Shigella* invasion.**

HeLa cells were challenged with bacteria for 30 min at 37°C, fixed and processed for immunofluorescence staining. Merge: maximal projections of confocal planes. Red, bacterial LPS; green, vinculin; blue, F-actin. vinculin: vinculin labeling. **a**, **b**, representative micrographs. *ipaA* mutant complemented with: full-length IpaA (WT); control vector (*ipaA*); IpaAΔVBS1-2 K498E (K498E). Scale bar = 5 µm. **a**, ROI delineated by: solid lines, actin foci (F); dotted lines, bacterial bodies (b). Vinculin recruitment at the bacterial body was quantified as the ratio of the average fluorescence intensity of vinculin labeling of (b) over that of (F). **b**, HV+ FA: large vinculin adhesions were quantified from images corresponding to the confocal plane of the cell basal surface as described in the Star Methods section.

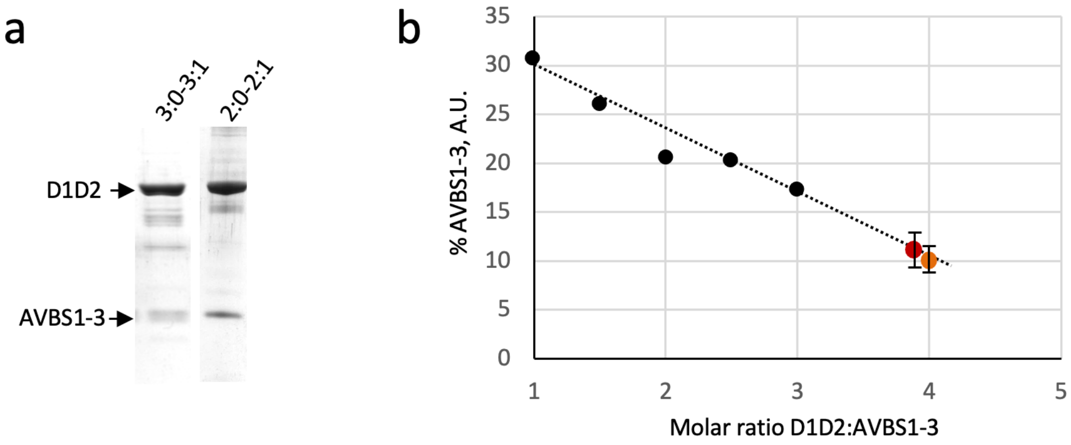

**Supplementary Fig. 2. Determination of the D1D2:AVBS1-3 molar ratio in peaks obtained in SEC-MALS analysis.** **a**, Representative SDS-PAGE analysis using a 10% polyacrylamide gel followed by Coomassie blue staining of the fractions corresponding to the 3:0-3:1 and 2:0-2:1 peaks shown in the SEC-MALS in Fig. 2f. The arrows point at D1D2 and AVBS1-3. **b**, the band integrated intensities were quantified using ImageJ. The values obtained for AVBS1-3 were multiplied by a factor of 3.15 to account for the difference in mass relative to D1D2. The percent of AVBS1-3 relative to the D1D2 and AVBS1-3 total amounts (% AVBS1-3) was plotted as a function of the D1D2:AVBS1-3 molar ratio. The values correspond to the average determination $\pm$ SEM (N = 3). Black circles: reference samples using defined D1D2:AVBS1-3 molar ratio. Red circles: molar ratio determined for the indicated peak fraction. Red: 3:0-3:1 peak. Orange: 2:0-2:1 peak. The determined D1D2:AVBS1-3 molar ratio of 3.8 and 3.9 for the 3:0-3:1 and 2:0-2:1 peaks, respectively, are consistent with a mixture of hetero- and homo-oligomers (2:0-2:1 and 3:0-3:1) in each peak.

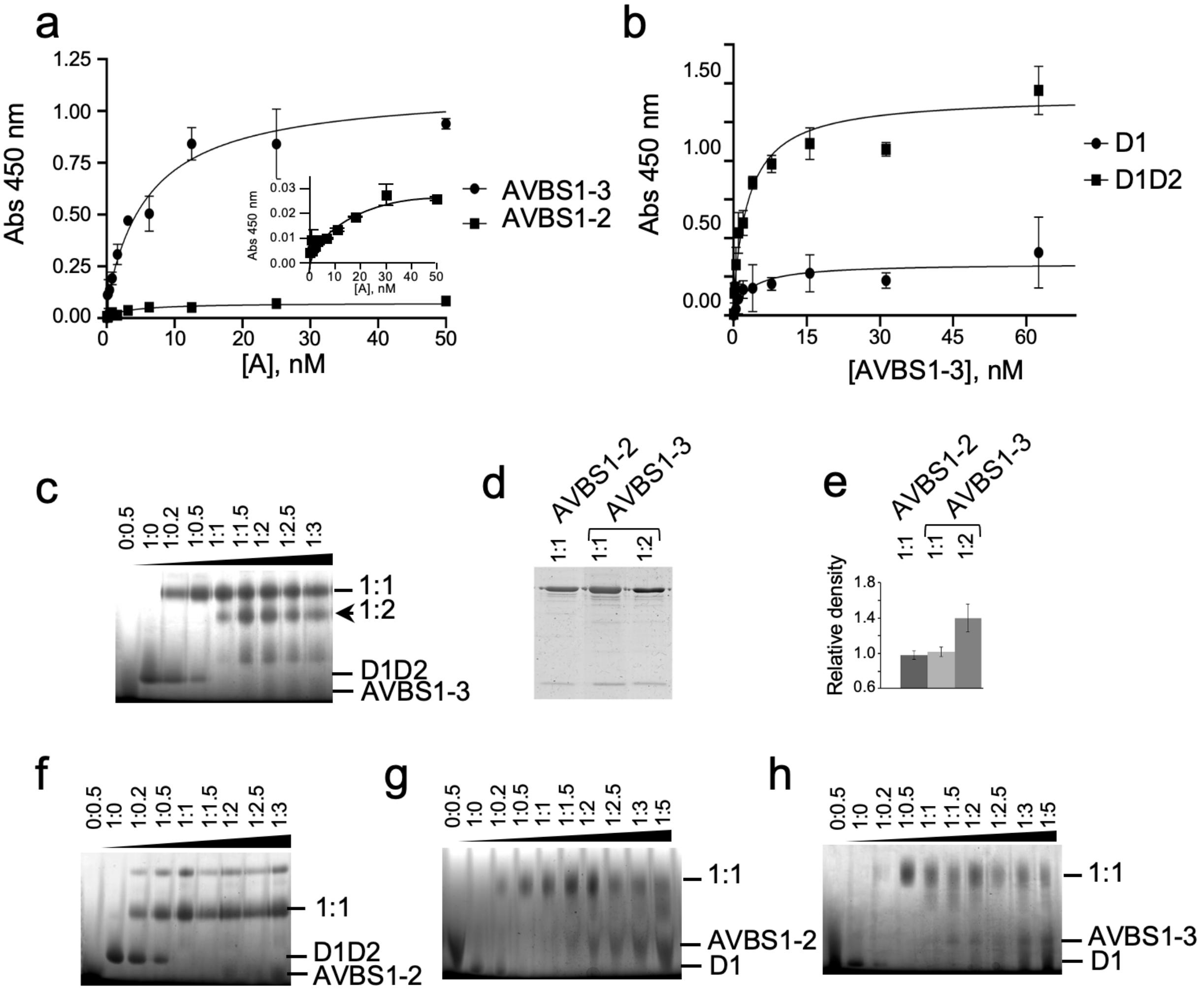

**Supplementary Fig. 3. IpaA VBS3 reveals multiple binding sites on vinculin.**

**a, b,** Solid phase binding assays. **a,** coating: vinculin; ligands: AVBS1-3 (solid circles); AVBS1-2 (solid squares). The inset shows binding of AVBS1-2 with an extended Y-axis. **b,** coating: D1 (solid circles) or vinculin D1D2 (solid squares); ligand: AVBS1-3. **c, f-h,** BN-PAGE in 6-18% polyacrylamide gradient gels and Coomassie staining analysis of D1D2:AVBS1-3 (**c**), D1D2:AVBS1-2 (**f**), D1:AVBS1-2 (**g**) or D1:AVBS1-3 (**h**) complexes. The molar ratio is indicated above each lane. Arrowheads indicate protein alone, or complex migration at the indicated molar ratio. **d**, bands were recovered from BN-PAGE and analyzed in a second dimension SDS-PAGE in a 15% poly-acrylamide gel and Coomassie staining. Bands were analyzed by densitometry. **e**, ratio of density values for the D1D2:AVBS1-2 complex (empty bar) and

D1D2:AVBS1-3 complexes corresponding to the upper (light grey bar) or lower (dark grey bar) shifts.

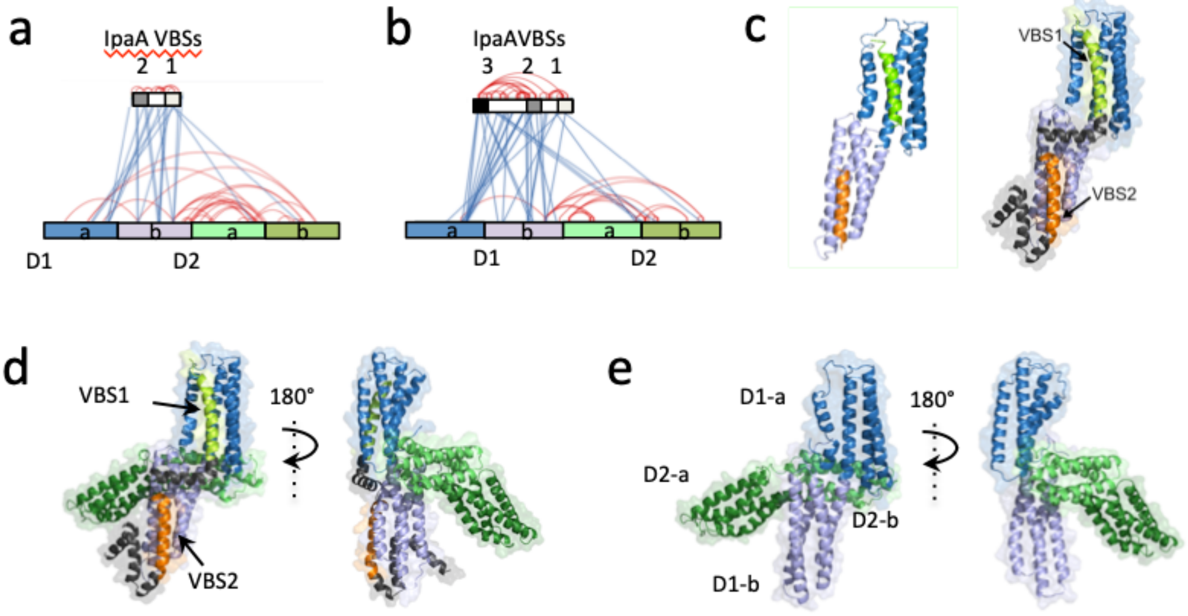

**Supplementary Figure 4. Structural models of vD1:AVBS1-2.**

**a**, **b**, EDC cross-link map from mass spectrometry analysis (LC-MS/MS) of vinculin D1D2-IpaA VBS1-2 (**a**) and D1D2-IpaA VBS1-3 (**b**) following extraction of 1:1 complexes from BN-PAGE. Blue lines: inter-molecular links. Red lines: intra-molecular links. Note the links between IpaA VBS3 and the D2 second bundle. Cross-linked residues are detailed in Suppl. Table 1. **c,** left, structure predicted from the resolved vD1: IpaA VBS1: and vD1:IpaA VBS2 crystal structures (Izard, Tran Van Nhieu et al. 2006, Tran Van Nhieu and Izard 2007)**;** right, structural model of vD1:IpaA VBS1-2. The model was established using RosettaCM protocol and accounts for 19 inter and intra-molecular cross-links out of 24 identified (Suppl. Table 1). Structural models of: **d**, D1D2:AVBS1-2. **e**, D1D2. AVBS1-2 were docked on the surface of Vinculin D1D2 using MS cross-link constraints. TX-MS protocol in combination with MS constraints was used to unify and adjust the final model, which justifies over 100 cross-links.

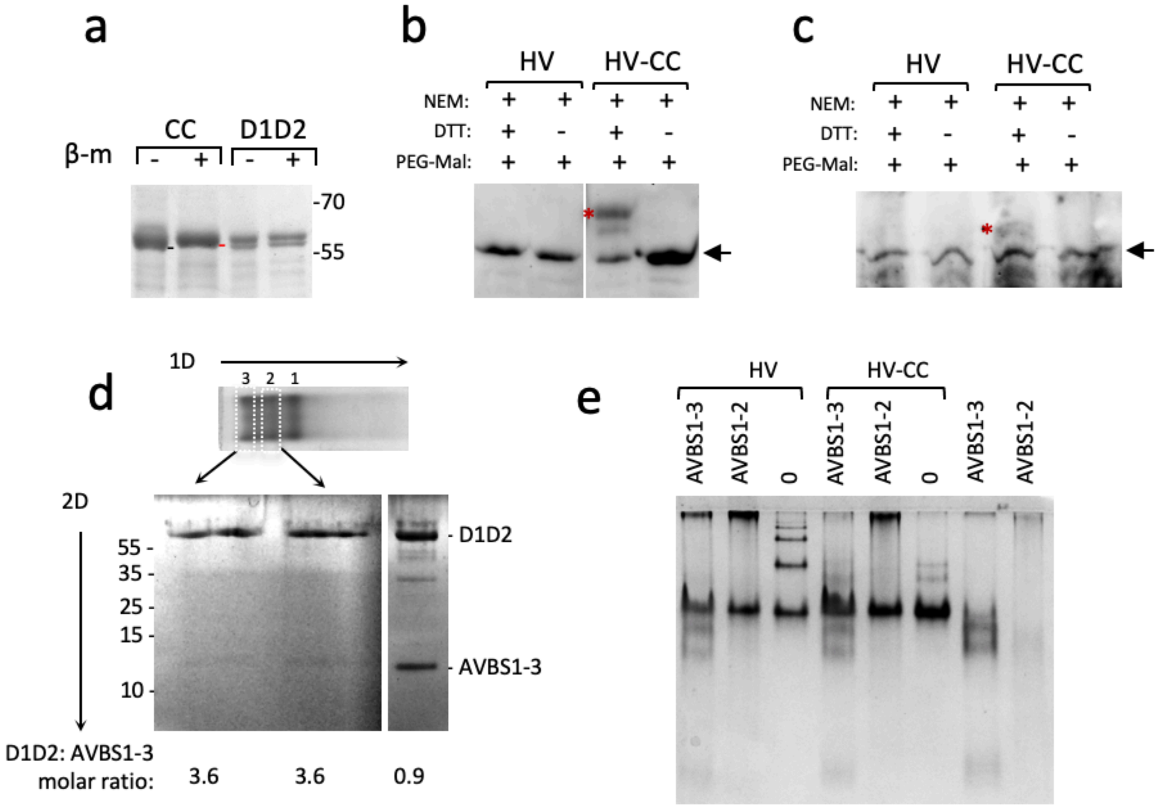

**Supplementary Figure 5. The Q68C A396C D1D2 mutant does not form high order complexes.** Representative gels revealed by Coomassie blue staining. **a**, Disulfide bridge formation in D1D2. D1D2: wild-type sequence. CC: D1D2 Q68C A396C. SDS-PAGE analysis using a 10 % polyacrylamide gel. + β-metOH: samples were boiled in Laemmli sample loading buffer containing 5 mM beta-mercaptoethanol prior to SDS-PAGE. The molecular weight markers in kDa are indicated. The black and red bars point to the respective migration of unreduced and reduced D1D2 Q68C A396C, respectively. **b, c,** Disulfide bridge formation in HV-CC. HV: full-length vinculin. HV-CC: HV Q68C A396C. Purified proteins (b) or samples pulled-down from lysates of HeLa cells transfected with pC1-HV8His or PC1-HVCC8His (Suppl. Star Methods) were treated with NEM, prior to reduction or not with DTT and treatment with PEG-Mal as indicated (Suppl. Star Methods). The red star points at the shifted band associated with the PEGylation of reduced disulfide bridge in HV-CC. **d**, bands corresponding to the shifts 2 and 3 observed upon incubation of D1D2 with AVBS1-3 (Fig. 3f) were dissected from native gels (1D). 2D: second dimension SDS-PAGE analysis using a 10 % polyacrylamide gel. Right strip: analysis of the reaction sample at a D1D2:AVBS1-3 molar ratio = 1. The numbers below the samples correspond to the D1D2:AVBS1-3 molar ratio determined by scanning densitometry. **e**, Clear Native-PAGE analysis in a 6-18% polyacrylamide gradient gel of HV and HV-CC complexes in the presence of AVBS1-3, AVBS1-2 at a molar ration of 1: 2 or buffer alone (0) as indicated.

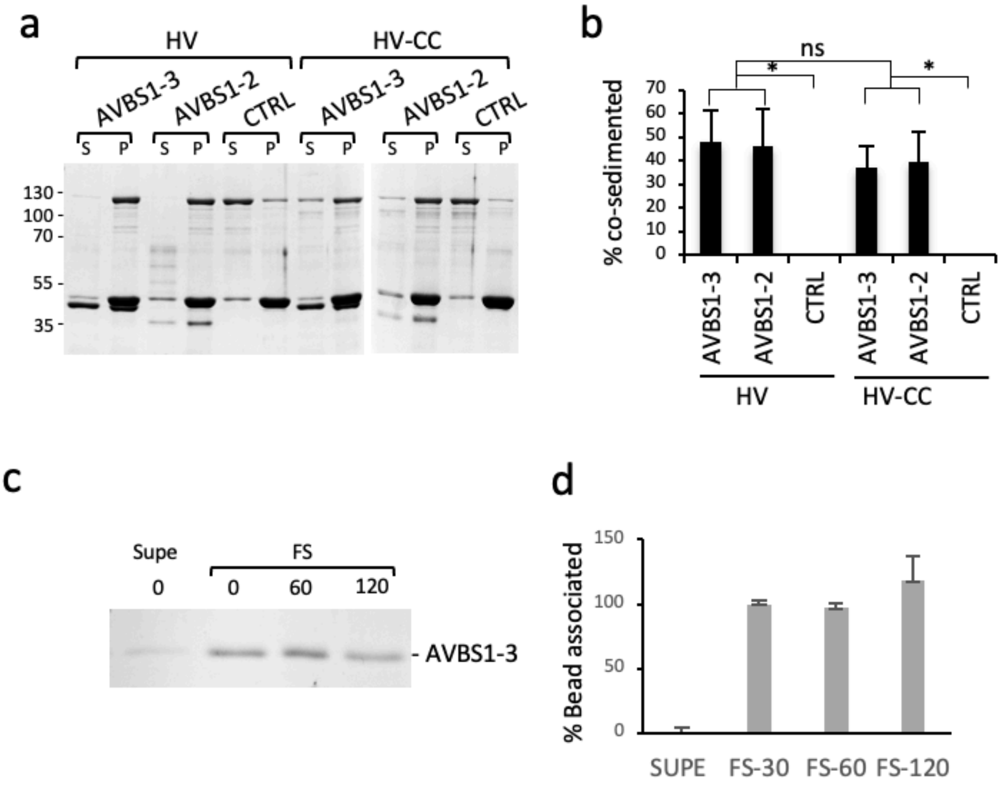

**Supplementary Figure 6. IpaA-induced vinculin oligomers mediate actin bundling at a distance from activation sites.**

**a, b,** actin sedimentation assays. HV: full-length human vinculin; HV-CC: cysteine-clamp derivative. Actin was allowed to polymerize at a final concentration of 10 μM in the presence of the indicated proteins. Samples were centrifuged at 110, 000 g for 30 min to pellet actin filaments. S: supernatant; P: pellet. **a**, **c**, representative SDS-PAGE analysis using a 10% polyacrylamide gel and Coomassie staining. **b**, **d**, the band integrated intensity corresponding to HV or HV-CC (**b**) or GST-AVBS1-3 (**d**) were quantified using ImageJ. Values are expressed as the percent of protein amounts in the pellet fraction relative to the total protein amounts in the supernatant and pellet fractions. **b**, percent of co-sedimented vinculin (n = 3, N = 3) or HV-CC (n = 3, N = 3). **c**, analysis of fluorosphere coating with GST-

AVBS1-3. Fluorospheres were coated with GST-AVBS1-3 at a final concentration of 2 μM in PBS, washed three times by centrifugation and resuspension in PBS, and incubated in F-actin buffer. At the time points indicated in min, samples were centrifuged for 2 min. Beads were resuspended in Laemmli loading sample buffer, incubated for 10 min at 95C and bead-associated protein were analyzed by SDS-PAGE using a 10% polyacrylamide gel. The supernatant was subjected to trichloroacetic acid precipitation at a final concentration of 5%, prior to acetone wash and SDS-PAGE analysis. Supe: supernatant. FS: beads-associated samples. **d**, average percent of the initial bead-associated amounts of GST-AVBS1-3. N= 3.

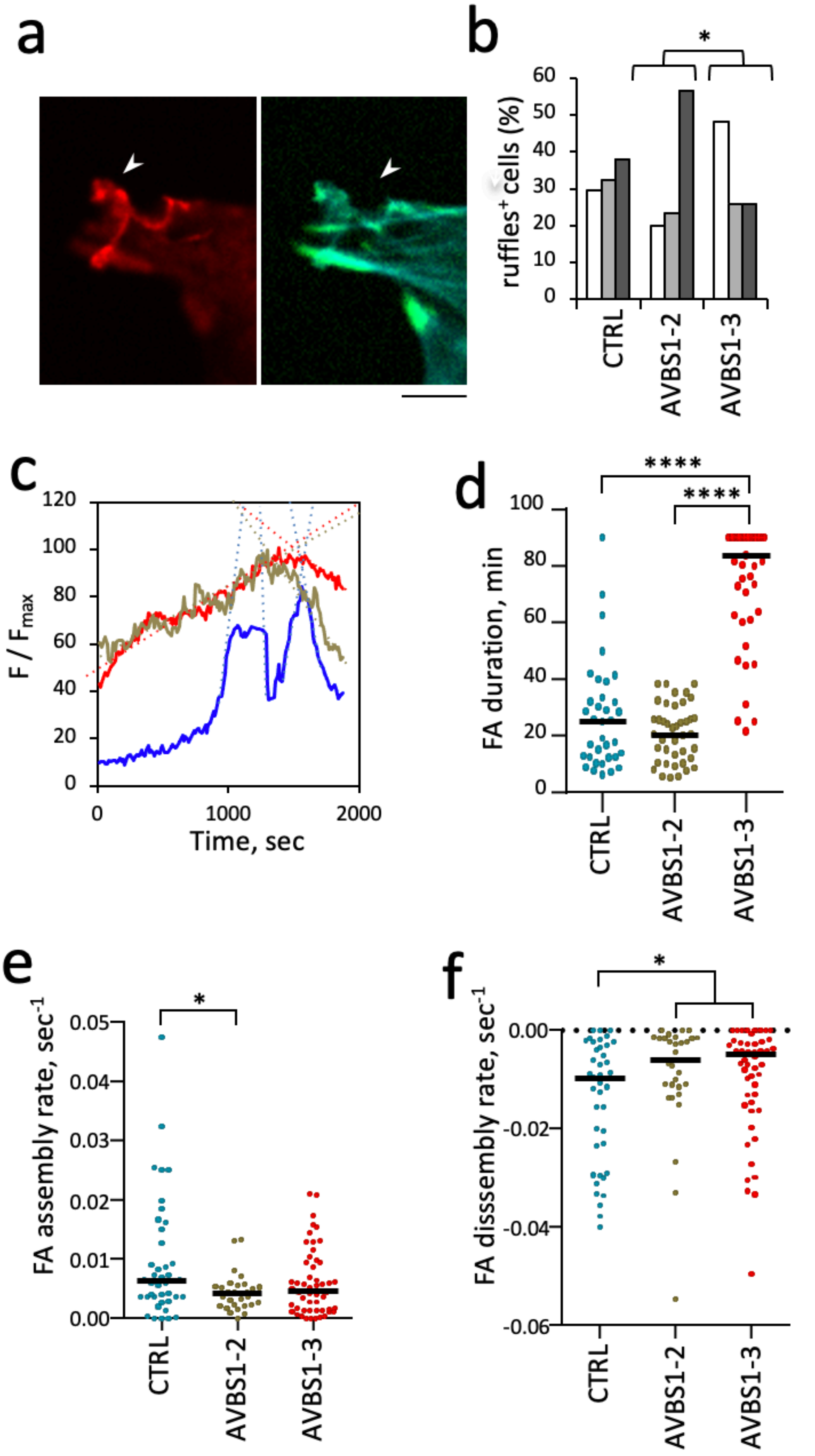

**Supplementary Figure 7. Actin ruffles and TIRF analysis of FA dynamics in AVBS1-2 and AVBS1-3 transfectants. a b**, CTRL: C2.7 cells. IpaA VBS1-2: GFP-IpaA VBS1-2 transfectants. IpaA VBS1-3:

GFP-IpaA VBS1-3 transfectants. **a,** representative fluorescence micrographs. Arrows: adhesions; arrowheads: ruffles. Green: GFP; red: vinculin; cyan: actin. **b**, percent of cells with ruffles ± SEM. Cells with: no ruffles (empty bars); small ruffles (light grey bars); large ruffles (dark grey bars). *: Pearson's Chi-squared test (N=3, n > 30, p = 0.036). **c-f,** C2. 7 cells were transfected with vinculin-mCherry (HV), vinculin-mCherry and GFP-IpaA VBS1-2 (AVBS1-2) or vinculin and GFP-IpaA VBS1-3 (AVBS1-3). The dynamics of vinculin-mCherry-labeled FAs were analyzed by TIRF microscopy. **c**, traces correspond to the variations of average fluorescence intensity of a representative single FA (F) normalized to its maximal average fluorescence intensity over the analyzed period in seconds ($F_{max}$). Blue: vinculin; green: vinculin + AVBS1-2; red: vinculin + AVBS1-3. **d**, FA duration. **e**, **f**, instant assembly (**b**) or disassembly (**c**) rates were inferred from the slopes of linear fits as depicted in a), with a Pearson correlation value > 0.85. HV: n = 41, N = 3; HV + AVBS1-2: n = 31, N = 2; HV + AVBS1-3: n = 55, N = 3. Mann-Whitney U test. *: $p < 0.05$.

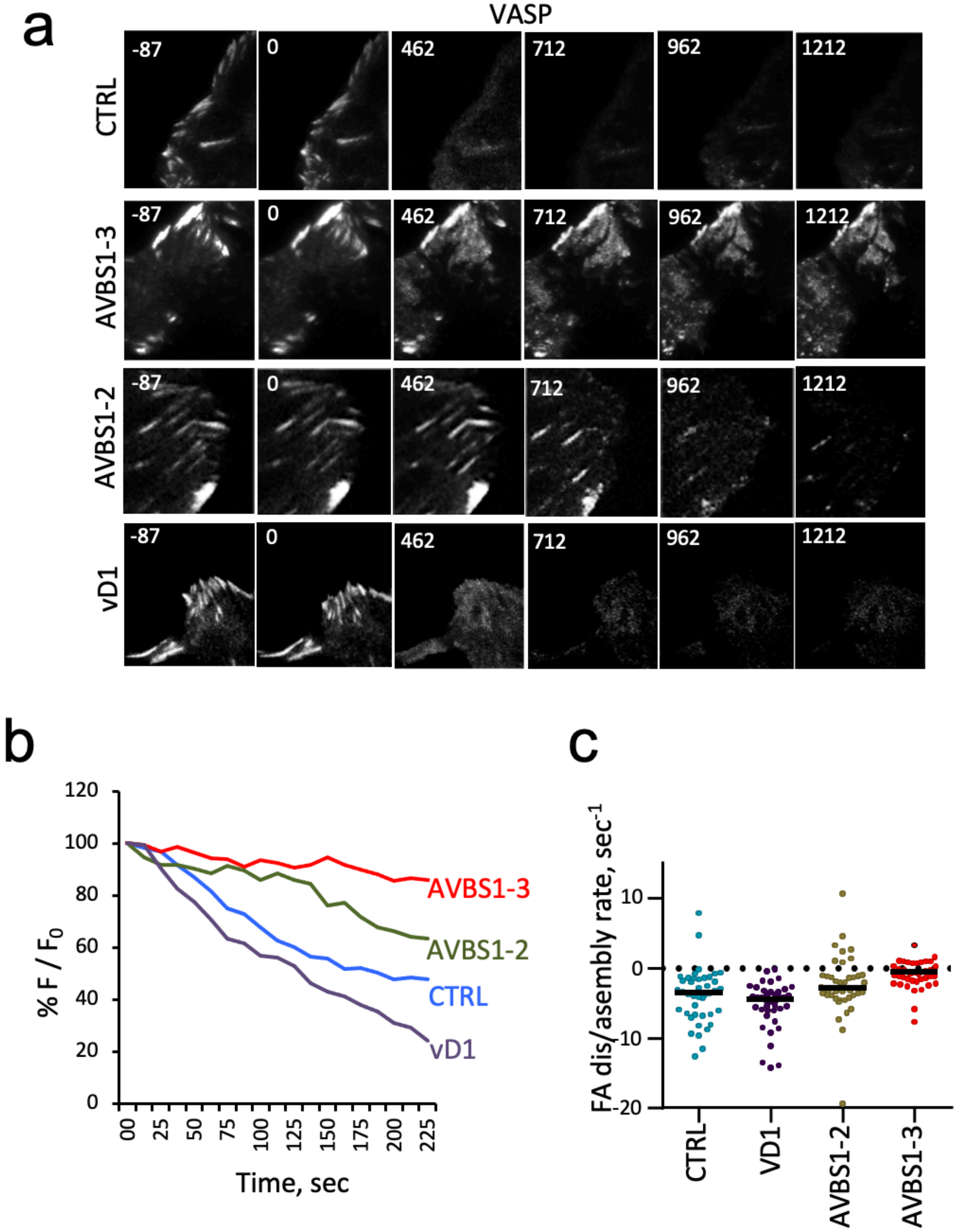

**Supplementary Figure 8. IpaA-mediated vinculin supra-activation stimulates cell adhesion independent of mechanotransduction.** TIRF microscopy of C2.7 cells transfected with mCherry-

VASP alone (CTRL) or co-transfected with GFP-AVBS1-2 (AVBS1-2), GFP-vD1 (vD1), or GFP-AVBS1-3 (AVBS1-3). Adhesion kinetic parameters were determined from time-lapse acquisitions following cell treatment with 100 µM Y-27632. **a,** representative time series acquisitions. Numbers indicated the elapsed time in seconds, with the inhibitor added at t = 0. Scale bar = 5 µm. **b**, % $F/F_0$: average fluorescence intensity of adhesions expressed as a percent of initial fluorescence. Solid lines: representative traces corresponding to single adhesions for the indicated samples in the corresponding color. The dashed redline illustrates FA assembly in AVBS1-3 transfected cell, seldom observed with the other samples. **c**, initial rates of adhesion assembly / disassembly inferred from linear fits. Number of adhesions analyzed: **c**, N = 4. CTRL: 42; IpaA VBS1-3: 43; IpaA VBS1-2: 40; vD1: 40. Dunn's multiple comparisons test. *: $p < 0.05$; ****: $p < 0.001$.

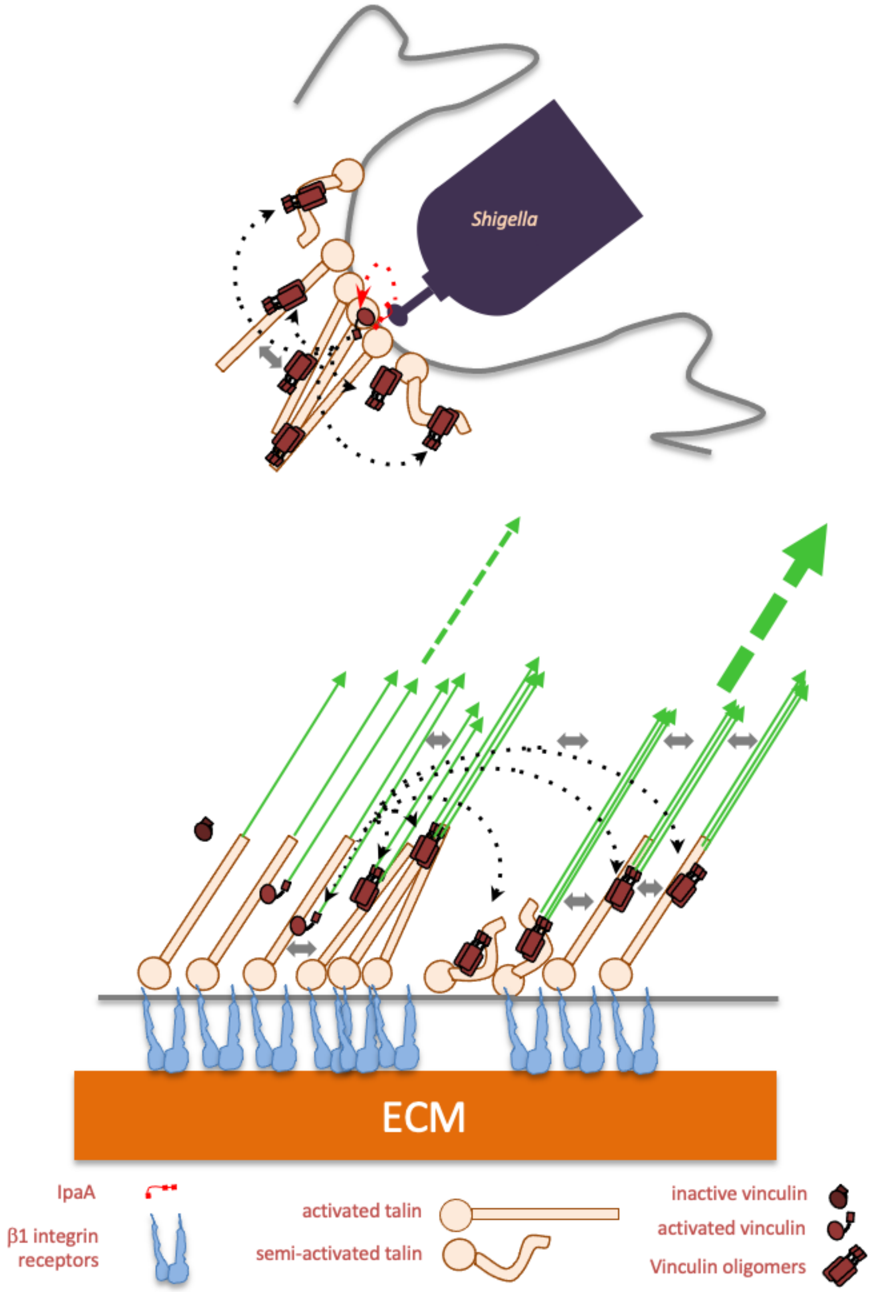

**Supplementary Figure 9. Scheme of IpaA-mediated vinculin supra-activation and oligomerization during *Shigella* invasion and FA maturation.** Top, IpaA induces vinculin supra-activation through the concerted action of the three IpaA VBS1-3 (red dotted arrow) at bacterial contact sites during *Shigella* invasion, leading to vinculin homo-oligomers diffusing away from activation sites (black dotted arrows). These vinculin oligomers are further clustered by talin, resulting in expansion of the bacterial adhesion structures. Bottom, vinculin canonical activation by talin VBSs combined with acto-myosin pulling force during mechanotransduction leads to supra-activation and oligomer formation. These oligomers diffuse from activation sites (black dotter arrows) and are captured by talin VBSs and actin fibers (green), leading to actin bundling and focal adhesion expansion associated with increased pulling force (dotted green arrows).

**SUPPLEMENTARY TABLES**

| # | Exp. MH$^{+}$ | Primary Score | Peptide Sequence | IpaA VBS1-2 | Hv D1 |
|---|---|---|---|---|---|
| 32670 | 4457.1 | 3.28 | IDDTSAELLTDDISDLKNNN**D**ITAENNNIYK - MA**K**MIDER | D604 | K173 |
| 32798 | 1924.16 | 2.61 | INN**K**LK - **E**LLPVLISAMK | K540 | E200 |
| 33042 | 4613.23 | 3.70 | IDDTSAELLT**D**DISDLKNNNDITAENNNIYK - NLGPGMT**K**MAK | D594 | K170 |
| 33042 | 4613.23 | 3.70 | IDDTSAELLTD**D**ISDLKNNNDITAENNNIYK - NLGPGMT**K**MAK | D595 | K170 |
| 33042 | 4613.23 | 4.11 | IDDTSAELLTDDISDLKNNN**D**ITAENNNIYK - NLGPGMT**K**MAK | D604 | K170 |
| 33322 | 5310.64 | 4.48 | IDDTSAELLTDDISDL**K**NNNDITAENNNIYK - VGKETVQTTE**D**QILKR | K600 | D67 |
| 33334 | 2347.28 | 2.51 | DVTTSLS**K**VLK - MSA**E**INEIIR | K625 | E240 |
| 33432 | 2138.26 | 2.88 | VL**K**NINKD - **E**LLPVLISAMK | K628 | E200 |
| 33496 | 2315.32 | 3.09 | AA**K**DVTTSLSK - **E**LLPVLISAMK | K617 | E200 |
| 34822 | 2642.55 | 2.68 | AKEVSSALS**K**VLSK - **E**LLPVLISAMK | K579 | E200 |
| 38218 | 3685.86 | 3.87 | NIN**K**D - TIESILEPVAQQISHLVIMHEEG**E**VDGK | K632 | E31 |

| | | | | | |
|---|---|---|---|---|---|
| 38218 | 3685.86 | 3.66 | NIN**K**D - TIESILEPVAQQISHLVIMH**E**EGEVDGK | K632 | E28 |
| **#** | **Exp. MH+** | **Primary Score** | **Peptide Sequence** | **Hv D1** | |
| 34075 | 2342.29 | 3 | **E**LLPVLISAMK - NLGPGMT**K**MAK | E200 | K170 |
| 36135 | 3088.67 | 2.62 | NFTVE**K**MSAEINEIIR - **E**LLPVLISAMK | K236 | E200 |
| **#** | **Exp. MH+** | **Primary Score** | **Peptide Sequence** | **IpaA VBS1-2** | |
| 14281 | 3051.56 | 3.52 | NYVT**E**TNADTIDKNHAIYEK - INN**K**LK | E554 | K540 |
| 15477 | 3342.67 | 2.74 | NYVT**E**TNADTIDKNHAIYEK - A**K**EVSSALSK | E554 | K571 |
| 17603 | 3141.54 | 2.71 | NYVTETNADTID**K**NHAIYEK - **E**VSSALSK | K562 | K572 |
| 31359 | 4483.2 | 2.64 | IDDTSA**E**LLTDDISDLKNNNDITAENNNIYK - A**K**EVSSALSK | E590 | K571 |
| 31996 | 2965.5 | 2.94 | IDDTSAELLTDDIS**D**LK - AA**K**DVTTSLSK | D598 | K617 |
| 31996 | 2965.5 | 2.51 | IDDTSAELLT**D**DISDLK - AA**K**DVTTSLSK | D594 | K617 |
| 32705 | 4586.24 | 4.1 | IDDTSAELLTDDISDLKNNNDITA**E**NNNIYK - AA**K**DVTTSLSK | E608 | K617 |
| 33253 | 3291.74 | 3.81 | IDDTSAELLTD**D**ISDLK - AKEVSSALS**K**VLSK | D595 | K579 |

| | | | | | |
|---|---|---|---|---|---|
| 33997 | 5328.54 | 4.76 | IDDTSAELLTDDISDL**K**NNNDITAENNNIYK - ID**D**TSAELLTDDISDLK | K600 | D586 |
| 34007 | 4655.33 | 4.78 | IDDTSAELLT**D**DISDLKNNNDITAENNNIYK - DVTTSLS**K**VLK | D594 | K625 |

**Suppl. Table 1**. Cross-linked residues characterized in the D1:IpaA VBS1-2 complex (XL-amino acids of each protein are bolded in the sequences).

| **#** | **Exp. $MH^+$** | **Primary Score** | **Peptide Sequence** | **IpaA VBS1-2** | **Hv D1D2** |
|---|---|---|---|---|---|
| 20480 | 3470.68 | 3.86 | NYVT**E**TNADTIDKNHAIYEK - NLGPGMT**K**MAK | E554 | K170 |
| 23021 | 3883.92 | 3.71 | NYVTETNADTID**K**NHAIYEK - **E**TVQTTEDQILKR | K562 | E60 |
| 24707 | 3117.73 | 5.00 | AKEVSSALSKVLSK - VGKETVQTT**E**DQILK | K579 | E66 |
| 36724 | 4612.22 | 4.82 | IDDTSAELLTDDISDLKNNN**D**ITAENNNIYK - NLGPGMT**K**MAK | D604 | K170 |
| 39712 | 5255.49 | 3.79 | IDDTSAELLTDDIS**D**LKNNNDITAENNNIYK - NLGPGMTKMA**K**MIDER | D598 | K173 |
| 39893 | 5162.6 | 4.51 | IDDTSA**E**LLTDDISDLKNNNDITAENNNIYK - ALA**K**QVATALQNLQTK | E590 | K464 |

| | | | | | |
|---|---|---|---|---|---|
| 40011 | 2641.56 | 4.45 | AKEVSSALS**K**VLSK - **E**LLPVLISAMK | K579 | E200 |
| 41002 | 3047.64 | 3.99 | L**K**VTDANIR - GILSGTSDLLLTFD**E**AEVR | K542 | E128 |
| 41440 | 3944.01 | 3.86 | IDDTSAELLTDDIS**D**LK - AQQVSQGLDVLTA**K**VENAAR | D598 | K366 |
| 42699 | 3685.87 | 4.03 | NIN**K**D -TIESILEPVAQQISHLVIMHEEG**E**VDGK | K632 | E31 |
| **#** | **Exp. MH$^+$** | **Primary Score** | **Peptide Sequence** | **Hv D1D2** | |
| 25162 | 3323.71 | 4.54 | LNQA**K**GWLRDPSASPGDAGEQAIR - ALASI**D**SK | K281 | D274 |
| 30025 | 3169.64 | 4.03 | GQGSSPVAMQ**K**AQQVSQGLDVLTAK - V**E**NAAR | K352 | E368 |
| 30045 | 3278.66 | 3.69 | VLQLTSWDE**D**AWASK - **K**LEAMTNSKQSIAK | D275 | K381 |
| 30201 | 3695.87 | 3.82 | LNQA**K**GWLRDPSASPGDAGEQAIR - MSA**E**INEIIR | K281 | E240 |
| 31264 | 3251.77 | 5.03 | ALA**K**QVATALQNLQTKTNR - RQGKGDSP**E**AR | K464 | E458 |
| 31264 | 3251.77 | 4.50 | ALA**K**QVATALQNLQTKTNR - RQGKG**D**SPEAR | K464 | D455 |
| 31620 | 2516.36 | 3.44 | ALASIDS**K**LNQAK - MSA**E**INEIIR | K295 | E240 |
| 35256 | 3787.88 | 4.12 | GILSGTSDLLLTFD**E**AEVR - GSSHHHHHHSS**G**LVPR | E128 | G12 |
| 38115 | 3310.75 | 4.29 | GILSGTSDLLLTFD**E**AEVR - AVANSRPA**K**AAVH | E147 | K507 |

| | | | | | |
|---|---|---|---|---|---|
| 39508 | 3685.90 | 3.56 | GQGSSPVAMQ**K**AQQVSQGLDVLTAK - MSA**E**INEIIR | K352 | E240 |
| 40134 | 3129.61 | 2.19 | NQGIEEALKNRNFTVEKMSAEINEIIR | E224 | K236 |
| 40285 | 3184.71 | 4.81 | AQQVSQGLDVLTA**K**VENAAR - SLG**E**ISALTSK | K366 | E437 |
| 40296 | 2280.32 | 3.69 | **E**LLPVLISAMK - LNQA**K**GWLR | E200 | K281 |
| 40351 | 3255.72 | 4.00 | AQQVSQGLDVLTA**K**VENAAR - MSA**E**INEIIR | K366 | E240 |
| 40571 | 4536.51 | 4.44 | G**D**SPEARALAKQVATALQNLQTK - SLGEISALTS**K**LADLRRQGK | D455 | K444 |
| 40881 | 3704.92 | 4.32 | **K**IDAAQNWLADPNGGPEGEEQIR - **E**LLPVLISAMK | K387 | E200 |
| 40923 | 3100.63 | 3.94 | AQQVSQGLDVLTAKV**E**NAAR - **K**LEAMTNSK | E368 | K373 |
| 41473 | 3723.01 | 4.34 | GQGSSPVAMQ**K**AQQVSQGLDVLTAK - **E**LLPVLISAMK | K352 | E200 |
| 41482 | 4633.41 | 3.54 | TI**E**SILEPVAQQISHLVIMHEEGEVDGK - **K**LEAMTNSKQSIAK | E28 | K381 |
| 42201 | 3294.83 | 3.66 | AQQVSQGLDVLTA**K**VENAAR - **E**LLPVLISAMK | K366 | E200 |
| **#** | **Exp. MH$^+$** | **Primary Score** | **Peptide Sequence** | **IpaA VBS1-2** | |
| 15424 | 3050.56 | 3.57 | NYVT**E**TNADTIDKNHAIYEK - INN**K**LK | E554 | K540 |

| | | | | | |
|---|---|---|---|---|---|
| 16531 | 3340.67 | 3.55 | NYVTETNA**D**TIDKNHAIYEK - A**K**EVSSALSK | E558 | K571 |
| 16865 | 3341.67 | 4.11 | NYVT**E**TNADTIDKNHAIYEK - A**K**EVSSALSK | E554 | K571 |
| 34602 | 4584.25 | 4.05 | IDDTSA**E**LLTDDISDLKNNNDITAENNNIYK - AA**K**DVTTSLSK | E590 | K617 |
| 35202 | 4586.25 | 5.2 | IDDTSAELLTDDISDLKNNN**D**ITAENNNIYK - AA**K**DVTTSLSK | D594 | K617 |
| 35824 | 4583.26 | 3.43 | IDDTSAELLTDDISDLKNNNDITA**E**NNNIYK - AA**K**DVTTSLSK | E608 | K617 |
| 39340 | 4911.48 | 4.37 | IDDTSAELLTD**D**ISDLKNNNDITAENNNIYK - AKEVSSALS**K**VLSK | D595 | K579 |
| 39340 | 4911.48 | 4.06 | IDDTSAELLTDDIS**D**LKNNNDITAENNNIYK - AKEVSSALS**K**VLSK | D598 | K579 |
| 39340 | 4911.48 | 3.83 | IDDTSA**E**LLTDDISDLKNNNDITAENNNIYK - AKEVSSALS**K**VLSK | E590 | K579 |
| 39730 | 3291.74 | 3.64 | IDDTSAELLTD**D**ISDLK - A**K**EVSSALSKVLSK | D595 | K571 |
| 40244 | 4655.32 | 4.35 | IDDTSAELLT**D**DISDLKNNNDITAENNNIYK - DVTTSLS**K**VLK | D594 | K625 |
| 40244 | 4655.32 | 4.35 | IDDTSAELLTD**D**ISDLKNNNDITAENNNIYK - DVTTSLS**K**VLK | D595 | K625 |

**Suppl. Table 2**. Cross-linked residues characterized in the D1D2:IpaA VBS1-2 complex (XL-amino acids of each protein are bolded in the sequences).

| # | Exp. MH+ | Primary Score | Peptide Sequence | IpaA VBS1-3 | Hv D1D2 |
|---|---|---|---|---|---|
| 13237 | 3014.59 | 3.29 | DIT**K**STTEHR - VGK**E**TVQTTEDQILKR | K530 | E66 |
| 14671 | 2518.38 | 2.49 | SKDIT**K** - VGK**E**TVQTTEDQILKR | K526 | E66 |
| 14791 | 2556.44 | 2.35 | INN**K**LK - VGK**E**TVQTTEDQILKR | K540 | E66 |
| 17806 | 2621.29 | 3.10 | **G**SPGIPGDTYLTR - QQ**E**LTHQEHR | G517 | E181 |
| 20039 | 2571.40 | 3.08 | L**K**VTDANIR **- E**TVQTTEDQILKR | K542 | E66 |
| 20526 | 3651.79 | 3.37 | NYVT**E**TNADTIDKNHAIYEK - NS**K**NQGIEEALK | E554 | K219 |
| 20985 | 3469.67 | 3.01 | NYVTETNA**D**TIDKNHAIYEK - NLGPGMT**K**MAK | D558 | K170 |
| 22936 | 3468.66 | 2.62 | NYVT**E**TNADTIDKNHAIYEK - NLGPGMT**K**MAK | E554 | K170 |
| 23137 | 4011 | 3.29 | NYVTETNADTI**D**KNHAIYEK - VG**K**ETVQTTEDQILK | D561 | K59 |
| 25418 | 3725.82 | 3.27 | NYVTETNADTID**K**NHAIYEK - ETVQTT**E**DQILK | K562 | E66 |
| 25418 | 3725.82 | 3.72 | NYVTETNADTID**K**NHAIYEK - **E**TVQTTEDQILK | K562 | E60 |

| | | | | | |
|---|---|---|---|---|---|
| 27618 | 3161.66 | 3.21 | **G**SPGIPGDTYLTR - VGKETVQTT**E**DQILKR | G517 | E60 |
| 27618 | 3161.66 | 3.61 | **G**SPGIPGDTYLTR - VGKETVQTTE**D**QILKR | G517 | D67 |
| 28366 | 2918.61 | 3.18 | EVSSALS**K**VLSK - VGK**E**TVQTTEDQILK - | K579 | E66 |
| 29996 | 1977.96 | 3.06 | **G**SPGIPGDTYLTR - MI**D**ER | G517 | D176 |
| 36497 | 2772.45 | 3.78 | **G**SPGIPGDTYLTR - AQQVSQGL**D**VLTAK | G517 | D361 |
| 36615 | 4613.23 | 3.68 | IDDTSAELLT**D**DISDLKNNNDITAENNNIYK - NLGPGMT**K**MAK | D594 | K170 |
| 36632 | 2214.27 | 3.22 | A**K**EVSSALSK **-** **E**LLPVLISAMK - | K571 | E200 |
| 37536 | 3824.86 | 2.64 | **G**SPGIPGDTYLTR - KI**D**AAQNWLADPNGGPEGEEQIR | G517 | D389 |
| 40263 | 3413.78 | 3.79 | GSPGIPG**D**TYLTR - AQQVSQGLDVLTA**K**VENAAR | D484 | K366 |
| 40432 | 5992.94 | 5.1 | IDDTSA**E**LLTDDISDLKNNNDITAENNNIYK - GQGSSPVAMQKAQQVSQGLDVLTA**K** | E590 | K362 |
| 40679 | 2708.42 | 3.25 | S**K**DITK - GILSGTSDLLLTFD**E**AEVR | K526 | E128 |
| 40994 | 3074.69 | 3.12 | **K**VTNSLSNLISLIGTK - ETVQTTEDQILK | K498 | E66 |
| 41187 | 4612.22 | 2.86 | IDDTSA**E**LLTDDISDLKNNNDITAENNNIYK - NLGPGMT**K**MAK | E590 | K170 |
| 41816 | 2387.41 | 3.65 | DVTTSLS**K**VLK - **E**LLPVLISAMK | K625 | E200 |
| 43326 | 3137.65 | 3.41 | AAKDVTTSLS**K** - GILSGTSDLLLTFD**E**AEVR | K625 | E128 |

| | | | | | |
|---|---|---|---|---|---|
| 43537 | 3542.88 | 3.63 | IDDTSAELLT**D**DISDLK - ALA**K**QVATALQNLQTK | D594 | K464 |
| 43612 | 3604.91 | 3.34 | VTNSLSNLISLIGT**K**SGTQER - ETVQTT**E**DQILK | K513 | E66 |
| 45081 | 3350.71 | 3.48 | **G**SPGIPGDTYLTR - GILSGTS**D**LLLTFDEAEVR | G517 | D121 |
| 53474 | 5745.03 | 4.09 | NIN**K**D - TIESILEPVAQQISHLVIMHEEG**E**VDGKAIPDLTAPVAAVQAAVSNLVR | K632 | E31 |
| **#** | **Exp. $MH^+$** | **Primary Score** | **Peptide Sequence** | **Hv D1D2** | |
| 17582 | 1670.89 | 3.51 | VG**K**ETVQTT**E**DQILK | K59 | E66 |
| 17582 | 1670.89 | 3.38 | VG**K**ETVQTTE**D**QILK | K59 | D67 |
| 26000 | 3322.71 | 4.03 | LNQA**K**GWLRDPSASPGDAGEQAIR - ALASI**D**SK | K281 | D374 |
| 30901 | 3169.64 | 3.79 | GQGSSPVAMQ**K**AQQVSQGLDVLTAK - V**E**NAAR | K352 | E368 |
| 31031 | 3695.87 | 3.81 | LNQA**K**GWLRDPSASPGDAGEQAIR - MSA**E**INEIIR | K281 | E240 |
| 35827 | 3665.8 | 3.75 | **K**IDAAQNWLADPNGGPEGEEQIR - MSA**E**INEIIR | K387 | E240 |
| 38215 | 3040.71 | 3.43 | VG**K**ETVQTTEDQILKR - ELLPVLISAM**K** | K59 | K200 |
| 39734 | 3614.9 | 4.15 | GQGSSPVAMQ**K**AQQVSQGLDVLTAK - SLG**E**ISALTSK | K352 | E437 |
| 40218 | 3580.72 | 3.88 | VLQLTSWDEDAWAS**K**DTEAMK - MSA**E**INEIIR | K261 | E240 |

| | | | | | |
|---|---|---|---|---|---|
| 40418 | 2881.63 | 3.77 | QVATALQNLQT**K**TNR - ELLPVLISAM**K** | K476 | E200 |
| 40559 | 3184.71 | 4.71 | AQQVSQGLDVLTA**K**VENAAR - SLG**E**ISALTSK | K366 | E437 |
| 40560 | 2280.32 | 3.35 | ELLPVLISAM**K** - LNQA**K**GWLR | E200 | K281 |
| 40604 | 2553.46 | 3.78 | ALASIDS**K**LNQAK - ELLPVLISAM**K** | K276 | E200 |
| 41140 | 2892.69 | 3.97 | ALA**K**QVATALQNLQTK - ELLPVLISAM**K** | K464 | E200 |
| 41598 | 4173.16 | 5.49 | GQGSSPVAMQKAQQVSQGLDVLTA**K**VENAAR - KL**E**AMTNSK | K366 | E375 |
| 41598 | 4173.16 | 5.8 | GQGSSPVAMQKAQQVSQGLDVLTA**K**VENAAR - **K**LEAMTNSK | E368 | K373 |
| 41598 | 4173.16 | 4.98 | GQGSSPVAMQKAQQVSQGL**D**VLTAK - VENAAR**K**LEAMTNSK | D361 | K373 |
| 41682 | 3726.02 | 3.87 | GQGSSPVAMQ**K**AQQVSQGLDVLTAK - ELLPVLISAM**K** | K352 | E200 |
| 42387 | 3293.83 | 4.63 | AQQVSQGLDVLTA**K**VENAAR - ELLPVLISAM**K** | K366 | E200 |
| 42387 | 3293.83 | 3.3 | AQQVSQGLDVLTAKV**E**NAARK - LNQA**K**GWLR | E368 | K281 |
| **#** | **Exp. MH⁺** | **Primary Score** | **Peptide Sequence** | **IpaA VBS1-3** | |
| 17398 | 3339.66 | 3.31 | NYVTETNADTI**D**KNHAIYEK - A**K**EVSSALSK | D561 | K571 |
| 17630 | 3341.68 | 3.44 | NYVTETNA**D**TIDKNHAIYEK - A**K**EVSSALSK | D558 | K571 |

| | | | | | |
|---|---|---|---|---|---|
| 17630 | 3341.68 | 4.19 | NYVT**E**TNADTIDKNHAIYEK - A**K**EVSSALSK | E554 | K571 |
| 20071 | 3141.53 | 3.53 | NYVTETNADTID**K**NHAIYEK - AK**E**VSSALSK | K562 | E572 |
| 20150 | 3349.69 | 4.44 | NYVTETNA**D**TIDKNHAIYEK - L**K**VTDANIR | D558 | K542 |
| 20150 | 3349.69 | 4.03 | NYVTETNADTI**D**KNHAIYEK - L**K**VTDANIR | D561 | K542 |
| 35163 | 4584.24 | 3.59 | IDDTSAELLT**D**DISDLKNNNDITAENNNIYK - AA**K**DVTTSLSK | D594 | K617 |
| 35163 | 4584.24 | 3.49 | IDDTSAELLTD**D**ISDLKNNNDITAENNNIYK - AA**K**DVTTSLSK | D595 | K617 |
| 35382 | 4586.24 | 3.35 | IDDTSA**E**LLTDDISDLKNNNDITAENNNIYK - AA**K**DVTTSLSK | E590 | K617 |
| 35967 | 4584.24 | 4.42 | IDDTSAELLTDDIS**D**LKNNNDITAENNNIYK - AA**K**DVTTSLSK | D598 | K617 |
| 35967 | 4584.24 | 4.95 | IDDTSAELLTDDISDLKNNNDITA**E**NNNIYK - AA**K**DVTTSLSK | E608 | K617 |
| 35967 | 4584.24 | 4.52 | IDDTSAELLTDDISDL**K**NNNDITAENNNIYK - AAK**D**VTTSLSK | K600 | D618 |
| 35967 | 4584.24 | 4.22 | IDDTSAELLTDDISDLKNNNDITAENNNIY**K -** AAK**D**VTTSLSK | K614 | D618 |
| 35967 | 4584.24 | 4.94 | IDDTSAELLTDDISDLKNNN**D**ITAENNNIYK - AA**K**DVTTSLSK | D604 | K617 |

| | | | | | |
|---|---|---|---|---|---|
| 39881 | 2845.52 | 4.73 | VTNSLSNLISLIGT**K**SGTQER - **E**LQEK | K513 | E520 |
| 39881 | 2845.52 | 4.62 | VTNSLSNLISLIGT**K**SGTQER - ELQ**E**K | K513 | E523 |
| 40044 | 4655.33 | 5.07 | IDDTSAELLT**D**DISDLKNNNDITAENNNIYK - DVTTSLSKVL**K** | D594 | K625 |
| 40044 | 4655.33 | 4.86 | IDDTSAELLTD**D**ISDLKNNNDITAENNNIYK - DVTTSLSKVL**K** | D595 | K625 |
| 40044 | 4655.33 | 4.45 | IDDTSAELLTDDIS**D**LKNNNDITAENNNIYK - DVTTSLSKVL**K** | D598 | K625 |
| 40044 | 4655.33 | 4.44 | IDDTSA**E**LLTDDISDLKNNNDITAENNNIYK - DVTTSLSKVL**K** | E590 | K625 |
| 40065 | 3092.6 | 3.51 | IDDTSAELLT**D**DISDLK - EVSSALSKVLS**K** | D594 | K579 |
| 40065 | 3092.6 | 3.38 | IDDTSAELLTD**D**ISDLK - EVSSALSKVLS**K** | D594 | K579 |
| 40069 | 3093.61 | 3.36 | IDDTSAELLTDDIS**D**LK - EVSSALSKVLS**K** | D598 | K579 |
| 40095 | 2988.61 | 4.48 | VTNSLSNLISLIGT**K**SGTQER - VT**D**ANIR | K513 | D545 |
| 40448 | 3252.74 | 3.81 | VTNSLSNLISLIGT**K**SGTQER - ETIF**E**ASKK | K513 | E494 |
| 40603 | 5153.62 | 4.36 | IDDTSAELLT**D**DISDLKNNNDITAENNNIYK - **K**VTNSLSNLISLIGTK | D594 | K498 |
| 40603 | 5153.62 | 4.46 | IDDTSAELLTD**D**ISDLKNNNDITAENNNIYK - **K**VTNSLSNLISLIGTK | D595 | K498 |

| | | | | | |
|---|---|---|---|---|---|
| 40603 | 5153.62 | 4.56 | IDDTSAELLTDDIS**D**LKNNNDITAENNNIYK - **K**VTNSLSNLISLIGTK | D598 | K498 |
| 40603 | 5153.62 | 3.99 | IDDTSAELLTDDISDLKNNN**D**ITAENNNIYK - **K**VTNSLSNLISLIGTK | D604 | K498 |
| 40606 | 5153.62 | 3.53 | IDDTSA**E**LLTDDISDLKNNNDITAENNNIYK - **K**VTNSLSNLISLIGTK | E590 | K498 |
| 41116 | 2593.44 | 3.85 | **K**VTNSLSNLISLIGTK - **E**TIFEASK | K498 | E490 |
| 41142 | 3909.09 | 4 | **E**TIFEASKKVTNSLSNLISLIGTK - **G**SPGIPGDTYLTR | E490 | G477 |

**Suppl. Table 3**. Cross-linked residues characterized in the D1D2 -IpaA VBS1-3 1:1 complex (XL-amino acids of each protein are bolded in the sequences)

### STAR METHODS

### RESOURCE AVAILABILITY

#### Lead contact

#### Materials availability

Availibility of plasmids, resources and reagents generated in this study may be subjected to restriction due to patent application N°PVT/ EP2016/073287.

#### Data and code availability

Any additional information required to reanalyze the data reported in this paper is available from the lead contact upon request.

#### Data

All data reported in this paper not related to the patent application N°PVT/ EP2016/073287 will be shared by the lead contact upon request.

#### Code

This paper does not report original code.

### METHODS DETAILS

#### Plasmids and constructs

Human vinculin constructs were generated by polymerase chain reaction using the forward primer 5’ GCGCATATGCCAGTGTTTCATACG-3’ and reverse primers 5’-CGTCGACTCACCAGGCATCTTCATCGGC-3’ for D1 (residues 1-258) or 5’-CGTCGACTCAGTGTACAGCTGCTTTG-3’ for D2 (residues 1-492) using a plasmid containing full-length octahistidine-tagged human vinculin (pET3a-vinculin 8His, residues 1–1,066), as template (Bakolitsa, Cohen et al. 2004), and cloned into the NdeI-SalI sites of pet15b (Novagen) to obtain pET15b-D1 and pET15b-D1D2, respectively. The Q68C and A396C cysteine substitution for the cysteine clamp were introduced into pet15b-D1D2 by site-directed mutagenesis using the 5’-GAGACTGTTCAAACCACTGAGGATTGCATTTTGAAG-3’ and 5’-ATCGATGCTGCTCAGAACTGGCTTTGCGATCCAAAT-3’ primers, respectively. The pGFP-vD1 plasmid was generated by polymerase chain reaction using the forward primer 5’-ACCCGGGATCCCGCC-3’ and reverse primer 5’-ACCCGGGACCAGGCA-3’, and cloned into pEGFP. The pmCherry-human vinculin (HV) and pmCherry-VASP plasmids were from Addgene. Stealth siRNA anti-human vinculin was from Invitrogen (reference number 1299001). The cysteine clamp was introduced in pmCherry N1-vinculin by exchanging the NheI-PspXI fragment with the corresponding XbaI-PspXI fragment of pET15b-D1D2 -Q68C A396C. Introduction of the cysteine clamp in full length vinculin was performed by swapping the *SexAI-BsrgI* fragment from pET15b-D1D2-Q68C-A396C and pET3a-vinculin 8His.

The IpaA constructs GFP-AVBS1-2 and GFP-AVBS1-3 were generated by polymerase chain reaction (PCR) and cloning into pcDNA3.1 NT-GFP Topo TA (Invitrogen) using the 5'-TCAAAGGACATTACAAAATCC-3' and 5’-GCGATATCATGGCCAGCAAAGG-3’ forward primers, respectively, and the 5’-GCGCGGCCGCTTAATCCTTATTGATATTC-3’ reverse primer. The GST-AVBS1-3 construct was generated by PCR using 5’-GGCGAATTCCCGGAGACACATATTTAACACG-3’ forward and 5’-GCCGTCGACTTAATCCTTATTGATATTCT-3’ reverse primers and cloning into the *EcoRI-SalI* ofpGEX-4T-2 (GE Lifesciences). pGST-AVBS1-2 was previously described (Ramarao, Le Clainche et al. 2007). The pGFP-vD1 plasmid was generated by polymerase chain reaction using the forward primer 5'-ACCCGGGATCCCGCC-3' and reverse primer 5'-ACCCGGGACCAGGCA-3', and cloned into pEGFP. The pmCherry-human vinculin (HV) and pmCherry-VASP plasmids were from Addgene. Stealth siRNA anti-human vinculin was from Invitrogen (reference number 1299001). All constructs were verified by DNA sequencing. Plasmids pC1-HV8His and pC1HV-CC8His were generated by replacing the 1.1 Kb EcoRV-NotI fragment of pmCherryN1-HV and pmCherryN1-HV-CC, respectively, by the 764 bp EcoRV-NotI fragment from pET3a-vinculin 8His.

**Cell lines and bacterial strains**

HeLa cells (ATCC CCL-2) were incubated in RPMI (Roswell Park Memorial Institute) medium containing 5% FCS (fetal calf serum, Gibco®) in an incubator with 5% $CO_2$. C2.7 myoblasts (Mitrossilis, Fouchard et al. 2009), MEF and MEF vinculin null cells (Humphries, Wang et al. 2007) were routinely grown in DMEM 1 g / L glucose containing 10 % FCS in a 37°C incubator containing 10 % $CO_2$. For transfection experiments, cells were seeded at 2.5 x $10^4$ cells in 25 mm-diameter coverslips. Cells were transfected with 3 µg of pGFP-AVBS1-2 or pGFP-AVBS1-3 plasmids with 6 µl JetPEI transfection reagent (Polyplus) for 16 hours following the manufacturer's recommendations. C2.7 mice myoblasts cells were fixed in PBS containing 3.7% paraformaldehyde for 20 min at 21°C and permeabilized with 0.1% Triton X-100 for 4 min at 21°C.

The wild type *Shigella flexneri*, isogenic mutants, and complemented *ipaA* mutant strains, as well as wild type *Shigella* expressing the AfaE adhesin were previously described (Izard, Tran Van Nhieu et al. 2006). Bacterial strains were cultured in trypticase soy broth (TCS) medium at 37°C. When specified, antibiotics were added at the following concentrations: carbenicillin 100 µg/ml, kanamycin 20 µg/ml.

**Cell challenge with *Shigella* strains**

HeLa cells seeded at 4 x $10^5$ cells in coverslip-containing 34 mm-diameter wells the day before the experiment. After 16 hours, cells were challenged with *Shigella* strains coated with poly-L-lysine, as follows. Bacteria grown to an $OD_{600\ nm}$ of 0.6 - 0.8 were washed three-times by successive centrifugation at 13 Kg for 30 sec and resuspension in EM buffer (120 mM NaCl, 7 mM KCl, 1.8 mM $CaCl_2$, 0.8 mM $MgCl_2$, 5 mM glucose, and 25 mM HEPES, pH = 7.3). Samples were resuspended in EM buffer containing 50 µg/ml poly-L-lysine and incubated for 15 min at 21°C, washed three times in EM buffer and resuspended in the same buffer at a final OD of $OD_{600\ nm}$ = 0.2. Cell samples were washed three times in EM buffer and challenged with 1 ml of the bacterial suspension and incubated at 37°C. Samples were fixed with PBS containing 3.7% PFA after 30 min incubation. Samples were processed for immunofluorescence microscopy.

**Immunofluorescence staining**

Cells were processed for immunofluorescence staining using the Vin11.5 anti-vinculin monoclonal antibody (ref. V4505, Sigma-Aldrich) and anti-mouse IgG antibody coupled to Alexa 546 (Jackson Research) and Phalloidin-Alexa 633 (Invitrogen), as described previously (Tran Van Nhieu and Izard 2007). Bacteria were labeled using anti-LPS rabbit polyclonal antibody followed by anti-rabbit IgG antibody coupled to Alexa 525 as described (Izard, Tran Van Nhieu et al. 2006). Samples were analyzed using an Eclipse Ti inverted microscope (Nikon) equipped with a 60 x objective, a CSU-X1 spinning disk confocal head (Yokogawa), and a Coolsnap HQ2 camera (Roper Scientific Instruments), controlled by the Metamorph 7.7 software. Analysis of fluorescent actin filaments was performed using a Leica confocal SP8 using a 63 x objective.

**Protein purification**

BL21 (DE3) chemically competent *E. coli* (Life Technologies) was transformed with the expression constructs. D1 and D1D2 were purified essentially as described (Papagrigoriou, Gingras et al. 2004, Park, Valencia-Gallardo et al. 2011). For the IpaA derivatives, bacteria grown until $OD_{600nm}$ = 1.0 were induced with 0.5 mM IPTG and incubated for another 3 hrs. Bacteria were pelleted and washed in binding buffer 25 mM Tris PH 7.4, 100 mM NaCl and 1 mM beta-mercaptoethanol, containing Complete™ protease inhibitor. Bacterial pellets were resuspended in 1/50th of the original culture volume and lyzed using a cell disruptor (One shot model, Constant System Inc.). Proteins were purified by affinity chromatography using a GSTrap HP affinity column (GE Healthcare) and size exclusion chromatography (HiLoad S200, GE Healthcare). Samples were stored aliquoted at -80°C at concentrations ranging from 1 to 10 mg/ml.

**Protein complex formation analysis**

Proteins were incubated at a concentration of 30 µM in binding buffer for 60 min at 4°C. Samples were analyzed by SEC-MALS using an HPLC (Shimadzu), a 24 ml Superdex 200 Increase 10/300 GL filtration column (GE Healthcare) and a MiniDAWN TREOS equipped with a refractometer Optilab T-rEX (Wyatt Technology) connected in series, to separate constructs according to their Stokes radius and determine the molar mass of macromolecules in solution. Data were analyzed using the ASTRA 6.1.7.17 software (Wyatt Technology Europe). Protein complex formation was visualized by PAGE under non-denaturing conditions using à 7.5% polycrylamide gel, followed by Coomassie blue staining.

**Solid-phase binding assay**

96–well Maxisorp (Nunc) ELISA plates were coated with 30 nM of full-length vinculin, vinculin constructs or IpaA proteins at the indicated concentrations in binding buffer (25 mM Tris PH 7.4, 100 mM NaCl and 1 mM β-mercaptoethanol). Samples were blocked with PBS-BSA 2%, washed and incubated with IpaA or vinculin proteins in binding buffer containing 0.2% BSA at room temperature for one hour. After incubation, the plates were washed and incubated with an anti-IpaA (dilution 1/2000$^{e}$) polyclonal primary antibody[3] or anti-vinculin (dilution 1/2000$^{e}$) Vin11.5 monoclonal antibody (Sigma-Aldrich) in binding buffer containing 0.2% BSA for one hour at room temperature. Plates were washed and incubated with an HRP-coupled secondary anti-rabbit or anti-mouse IgG antibody (1/32000$^{e}$) (Jackson ImmunoResearch) for one hour. The reaction was revealed by adding 100 µl of tetramethylbenzidine (Sigma-Aldrich) for 15 min, stopped by adding 50 µl of 0.66N $H_2SO_4$ and the absorbance was read at 450 nm (Dynatech MR400).

**BN-PAGE (Blue Native – Polyacrylamide Gel Electrophoresis) protein native gel analysis and complex cross-linking**

25 µM of vinculin constructs were incubated with different molar ratios of IpaA proteins in a 1X BN-PAGE buffer (250 mM ε-aminicaprionic acid and 25 mM Bis-Tris PH 7,0) at 4°C for one hour. The protein mixtures were separated in a one-dimension native BN-PAGE electrophoresis as described (Eubel and Millar 2009). For vinculin-IpaA protein ratio assay, vinculin-IpaA bands containing the complexes separated by BN-PAGE were cut, sliced and boiled in 2 x Laemmli SDS buffer followed by SDS-PAGE. The second dimension SDS-PAGE gels were stained (colloidal Coomassie staining) and the density of the bands was determined using Image J. The normalized vinculin:IpaA ratio of the complexes was compared using a non-parametric Kruskal-Wallis rank sum test (R statistical software).

For crosslinking vinculin-IpaA complex, bands containing the complexes were cut, sliced and electroeluted in native conditions (15 mM Bis-Tris pH 7.0 and 50 mM Tricine) inside a closed dialysis membrane (SpectraPor). The soluble complexes were recovered and their buffer exchanged twice into an amine-free cross-link buffer in 25 mM HEPES pH 7.0 containing 100 mM NaCl using 10MWCO ZEBA desalting columns (Thermo Scientific). The fractions containing the complexes were incubated for 1 hr at 4°C with 10 mM N-hydroxysulfosuccinimide and 5 mM EDC (Sigma-Aldrich) following the manufacturer's recommendations. The cross-linking reaction was stopped by adding 50 mM Tris pH 7.4 and incubating for 20 minutes. Samples were denaturated in 2x SDS Laemmli buffer for 5 min at 95°C and complexes were eluted from gel slices following SDS-PAGE.

**Liquid Chromatography Mass spectrometry (LC-MS)**

Complexes obtained after the cross-linking step were loaded onto a 4-20% polyacrylamide gradient gels and Coomassie stained. The bands containing the complexes were cut and submitted to tryptic digestion (Shevchenko, Tomas et al. 2006). The experiments were performed in duplicates for the 3 complexes D1:AVBS1-2, D1D2: AVBS1-2 and D1D2:AVBS1-3. Peptides obtained after tryptic digestion were analyzed on a Q Exactive Plus instrument (Thermo Fisher Scientific, Bremen) coupled with an EASY nLC 1 000 chromatography system (Thermo Fisher Scientific, Bremen). Sample was loaded on an in-house packed 50 cm nano-HPLC column (75 µm inner diameter) with C18 resin (1.9 µm particles, 100 Å pore size, Reprosil-Pur Basic C18-HD resin, Dr. Maisch GmbH, Ammerbuch-Entringen, Germany) and equilibrated in 98 % solvent A (H2O, 0.1 % FA) and 2 % solvent B (ACN, 0.1 % FA). A 120 minute-gradient of solvent B at 250 nL.min-1 flow rate was applied to separate peptides. The instrument method for the Q Exactive Plus was set up in DDA mode (Data Dependent Acquisition). After a survey scan in the Orbitrap (resolution 70 000), the 10 most intense precursor ions were selected for HCD fragmentation with a normalized collision energy set up to 28. Charge state screening was enabled, and precursors with unknown charge state or a charge state of 1 and >7 were excluded. Dynamic exclusion was enabled for 35 or 45 seconds respectively.

**Analysis of disulfide bridge by PEG-Maleimide modification**

The determination of disulfide bridge formation in HV-CC was performed as previously described with minor modifications (Braakman, Lamriben et al. 2017; Pant, Oh, and Mysore 2021). Briefly, for in vitro determination, 2 µg of purified HV or HV-CC were first incubated with 50 mM NEM PEG-Mal, NEM (N-ethylmaleimide), prior to reduction by incubation in 50 mM DTT (dithiothreitol). Alternatively, control samples were first reduced by incubation in 50 mM DTT, prior to NEM treatment. All samples were then incubated in 8 mM PEG-Mal (Sigma Aldrich, 63187). All incubation steps were carried out for 40 min at 21°C in 150 mM Tris pH 6.8, 0, 1%SDS (incubation buffer). Samples were precipitated and washed twice with acetone prior to resuspension between incubation steps. For determination of disulfide bridge formation in cells, HeLa cells were transfected with plasmids pC1-HV8His or pC1HV-CC8His the day preceding the experiment. Cells were scraped in 200 µls of ice-cold incubation buffer containing 1 mM AEBSF (4-(2-aminoethyl)benzenesulfonyl fluoride). Cell lysates were transferred to Eppendorf tubes on ice, 1800 uls of 150 mM Tris pH 6.8 containing 1 mM DTT (wash buffer) were added, and samples were clarified by centrifugation for 10 min at 13,000 g at 4°C. Clarified lysates were transferred to a fresh Eppendorf tube and subjected to pull-down by incubation with 40 µls of Ni Sepharose 6 Fast flow resin (Sigma Aldrich Pharmaceuticals) for 40 min at 4°C using an end-over-end roller. Samples were washed three times with 500 µls wash buffer. Samples were then subjected to NEM and DTT treatment prior to PEG-Mal modification as described above for purified proteins. Samples were resuspended in Laemmli loading sample buffer and SDS-PAGE analysis using a 7.5% polyacrylamide gel. For purified proteins, gels were analyzed by Coomassie blue staining. For proteins pulled-down from cell lysates, samples were analyzed by anti-vinculin Western blot analysis using the Vin11.5 mouse monoclonal antibody (Sigma Aldrich Pharmaceuticals) at a 1:5000 dilution.

**Actin sedimentation assays**

Actin was purified from rabbit muscle as described previously (Ciobanasu, Faivre et al. 2015). For actin co-sedimentation assays 5 µM actin was incubated in F-actin buffer (2 mM Tris pH7.5, 0.2 mM $CaCl_2$, 0.5 mM beta-mercaptoethanol, 0.2 mM ATP, 100 mM KCl , 2 mM $MgCl_2$) for 60 min at 21°C. When indicated, HV, HV-CC, AVBS1-3 and AVBS1-2 were added at a 2 µM final concentration. Samples were centrifuged at 110,000 g for 30 min at 4°C. For actin bundling assays, actin was allowed to polymerize at the indicated concentration and samples were centrifuged at 14,000 g for 10 min at 31°C. Proteins in pellets and supernatants were analyzed by SDS-PAGE on a 10% polyacrylamide gel followed by Coomassie blue staining. For quantification, protein band integrated densities were determined using ImageJ. The percentage of pelleted protein is calculated relative to the total amounts of the corresponding protein in the supernatant and pellet.

**Fluorescent microscopy analysis of actin bundling by vinculin oligomers.**

HV and HV-CC were labeled using BodipyTM FL NHS Ester (Succinimidyl Ester) (ThermoFisher, D2184) following the manufacturer's instruction.  Actin was fluorescently labeled with AlexaFluor 594 Succinimidyl Ester as previously described (Ciobanasu, Faivre et al. 2015). To immobilize protein on beads, 30 µls of 1 µm-diameter red Fluorospheres (Molecular Probes, F8887) were incubated with 2 µM GST-AVBS1-3, GST-AVBS1-2 or GST as a control in 200 µl PBS for 120 min with end-over-end rolling at 4°C. Beads were washed three-times by successive centrifugation for 2 min at 14,000 g and resuspension in 200 µl PBS. 100 nm-diameter red Fluorospheres were coated with 2 µM of talin H1-H4 (Papagrigoriou, Gingras et al. 2004) using a similar procedure. For each sample, 30 µl of the coated-bead suspension were centrifuged 2 min at 14,000 g, the supernatant was discarded and pelleted beads were resuspended in F-actin buffer containing 3.5 µM actin, 1.5 µM AlexaFluor 594 -labeled actin, and HV or HV-CC at 1 µM final concentration in a total volume of 10 µl. Samples were incubated for 15 min at 21°C, Phalloidin AlexaFluor 594 (Thermofisher, A12831) was added at a final concentration of 100 nM and incubation was allowed to proceed for another 45 min. Samples were mixed with DAKO mounting medium (DAKO, S3023), placed on a slide and covered by a 22 x 22 mm coverslip. Samples were analyzed using a Leica SP8 confocal microscope using a 63 x oil immersion objective, at an image resolution of 2048 x 2048, zoom 4.

**Data analysis**

The identification of cross-linked peptides from LC-MS data was performed using SIM-XL v. 1.3 (Lima, de Lima et al. 2015), with the following search parameters: EDC as cross-linker, a tolerance of 20 ppm for precursor and fragment ions, trypsin fully specific digestion with up to three missed cleavages. Carbamidomethylation of cysteines was considered as a fixed modification. All initial identification of cross-linked peptides required a primary score of SIM-XL greater than 2.5 for inter-links and 2.0 for intra-links or loop-links. As single incorrect cross-link identification might lead to a different model, a manual post-validation of the search engine results at the MS2 level was thus performed. A 2D-map showing the protein-protein interaction was generated as an output (Figs. 2a,b). Only peptides present in the 2 replicates are gathered in Supplementary Tables 1-3 and were used for the modeling.

**Modeling**

We used the distance constraints obtained from cross-linking MS data (Suppl. Tables 1-3) to guide the protein structure modeling using the TX-MS protocol as described by Hauri, Khakzad et al. (Hauri, Khakzad et al. 2019). In short, TX-MS uses the Rosetta comparative modeling protocol (RosettaCM) (Song, DiMaio et al. 2013), and the flexible backbone docking protocol (RosettaDock) (Gray 2006) to generate models and evaluate how well each model explains the MS constraints using a novel scoring function. Here, a total of 100,000 models was generated, of which the highest-scoring model is displayed in (Fig. 3c), supported by a total of 100 inter and intra-molecular cross-links.

**TIRF (Total Internal Reflection Microscopy) analysis**

C2.7 cells were transfected with pmCherry-vinculin or pmCherry-VASP and the indicated plasmids as described above. Samples were mounted onto a TIRF microscopy chamber on a stage of an Eclipse Ti inverted microscope (Nikon) equipped with an Apo TIRF 100 x N.A. 1.49 oil objective heated at 37°C. TIRF analysis was performed using the Roper ILAS module and an Evolve EM-CCD camera (Roper Scientific Instruments). When mentioned, Y-27632 was used at 100 µM. Image acquisition was performed every 12.5 seconds for 30 to 90 minutes.

**Invasion assays**

Tissue culture Transwell inserts (8 µm pore size; Falcon, Franklin Lakes, NJ) were coated for 3 hours with 10 µg of Matrigel following the manufacturer's instructions (Biocoat, BD Biosciences, San Jose, CA). Inserts were placed into 24-well dishes containing 500 µl of RPMI medium supplemented with 1% fetal calf serum. $5 \times 10^4$ melanoma cells were added to the upper chamber in 250 µl of serum-free RPMI medium. After 24 hours, transmigrated cells were scored by bright field microscopy. Experiments were performed at least three times, each with duplicate samples.

**QUANTIFICATION AND STATISTICAL ANALYSIS**

Vinculin clusters induced by AVBS1-2- and AVBS1-3-coated beads were analyzed using the ImageJ 2.1.0/1.53c software. Briefly, for each set of experiments, sum projection images of confocal planes were thresholded using identical parameters between samples. Clusters were detected using the “Analyze particle” plug-in, setting a minimal size of 570 $nm^2$. The vinculin cluster integrated fluorescence density was arbitrarily expressed as the percent of the mean integrated fluorescence density determined for vinculin clusters induced by GST-coated beads (Fig. 2) or AVBS1-2-coated beads (Fig.3).

Quantification of vinculin recruitment at the close vicinity of invading bacteria was performed on the sum projection of confocal planes corresponding to *Shigella*-induced actin foci. ROIs were drawn to delineate the actin foci (F) and the bacterial body (b) from the corresponding wavelength channel as shown in Fig. S1. The vinculin recruitment index was calculated as the ratio of the average fluorescence intensity corresponding to vinculin labeling associated with (b) corrected to background over that of (F). For the quantification of the number and size of large adhesion structures induced by *Shigella* invasion in HeLa cells, the confocal fluorescent microscopy plane corresponding to the vinculin-labeled cell basal plane was processed using the imageJ FFT / Bandpass followed by particle analysis plugins with a low size threshold set at 3.54 $µm^2$. A semi-automated protocol using Icy software was developed for the quantification of adhesion structures in C2.7 cells (de Chaumont, Dallongeville et al. 2012). Confocal fluorescent microscopy planes were used to detect vinculin structures using HK means thresholding and overlaid binary masks obtained from the threshold projections of F-actin labeled images (Max-entropy method). FAs were detected as spots positive for both vinculin mCherry and actin structures using Wavelet Spot Detector. The number of adhesions was analyzed using Dunn's multiple comparisons test. The statistical analysis of cell motility was performed in the R software. Medians were compared using a Wilcoxon rank sum test and dispersion by Median absolute dispersion (MAD) parameter.

**KEY RESOURCES TABLE**

| REAGENT or RESOURCE | SOURCE | IDENTIFIER |
|---|---|---|
| Antibodies | | |
| Vin11.5 | Sigma-Aldrich | ref. V4505 |
| Polyclonal anti-Shigella serotype V lipopolysaccharide | Valencia-Gallardo et al., 2019 | |
| Bacterial and virus strains | | |
| *Shigella flexneri* serotype V | Valencia-Gallardo et al., 2019 | M90T |
| *Shigella flexneri* serotype V *ipaA* mutant | Valencia-Gallardo et al., 2019 | M90T *ipaA* |
| Biological samples | | |
| Non applicable | | |
| Chemicals, peptides, and recombinant proteins | | |
| N-hydroxysulfosuccinimide | MERCK | 106627-54-7 |
| 1-ethyl-3-carbodiimide hydrochloride | Sigma-Aldrich | 25952-53-8 |
| N-ethylmaleimide | Sigma-Aldrich | 128-53-0 |
| dithiothreitol | Sigma-Aldrich | 16096-97-2 |
| PEG-Mal | Sigma-Aldrich | 63187 |
| BodipyTM FL NHS Ester | ThermoFisher | D2184 |

| | | |
|---|---|---|
| Y-27632 | Sigma-Aldrich | 129830-38-2 |
| Critical commercial assays | | |
| Non applicable | | |
| Deposited data | | |
| Non applicable | | |
| Experimental models: Cell lines | | |
| HeLa cells | ATCC | ATCC CCL-2 |
| C2.7 cells | Mitrossilis et al., 2009 | |
| MEF vinculin null cells | Humphries, Wang et al. 2007 | |
| Experimental models: Organisms/strains | | |
| Non applicable | | |
| Oligonucleotides | | |
| 5’ GCGCATATGCCAGTGTTTCATACG-3’ | This study | vD1 For |
| 5’-CGTCGACTCACCAGGCATCTTCATCGGC-3’ | This study | vD1 Rev |
| 5’-CGTCGACTCAGTGTACAGCTGCTTTG-3’ | This study | vD2 Rev |
| 5’-GAGACTGTTCAAACCACTGAGGATTGCATTTTGAAG-3’ | This study | HV-Q68C mut |
| 5’-ATCGATGCTGCTCAGAACTGGCTTTGCGATCCAAAT-3’ | This study | HV-A396C mut |
| 5’-ACCCGGGATCCCGCC-3’ | This study | GFP-vD1 For |
| 5’-ACCCGGGACCAGGCA-3’ | This study | GFP-vD1 Rev |
| 5'-TCAAAGGACATTACAAAATCC-3' | This study | GFP-AVBS1-2 For |
| 5’-GCGATATCATGGCCAGCAAAGG-3’ | This study | GFP-AVBS1-3 For |
| 5’-GCGCGGCCGCTTAATCCTTATTGATATTC-3’ | This study | GFP-AVBS Rev |
| 5’-GGCGAATTCCCGGAGACACATATTTAACACG-3’ | This study | GST-AVBS1-3 For |
| 5’-GCCGTCGACTTAATCCTTATTGATATTCT-3’ | This study | GST-AVBS1-3 Rev |
| 5’-ACCCGGGATCCCGCC-3’ | This study | GFP-vD1 For |

| | | |
|---|---|---|
| 5'-ACCCGGGACCAGGCA-3' | This study | GFP-vD1 Rev |
| Recombinant DNA | | |
| pET15b-D1 | This study | |
| pET15b-D1D2 | This study | |
| pET15b-D1D2-CC | This study | |
| pGFP-vD1 | This study | |
| pmCherry N1-HV-CC | This study | |
| pGFP-AVBS1-2 | This study | |
| pGFP-AVBS1-3 | This study | |
| pGST-AVBS1-2 | Ramaro et al., 2007 | |
| pGST-AVBS1-3 | This study | |
| pC1-HV8His | This study | |
| pC1HV-CC8His | This study | |
| pmCherry-human vinculin | Addgene | |
| pmCherry-VASP | Addgene | |
| Software and algorithms | | |
| Icy | De Chaumont et al., 2012 | |
| Rosetta modeling | Song et al., 2013 | |
| RosettaDock | Gray, 2006 | |
| ASTRA 6.1.7.17 | Wyatt Technology Europe | |
| Other | | |
| Non applicable | | |

**Suppl. movie 1.** TIRF analysis of vinculin-mCherry expressing C2.7 cells co-transfected with a GFP fusion to the indicated construct. The time is indicated in seconds.

**Suppl. movie 2.** TIRF analysis of vinculin-mCherry expressing C2.7 cells co-transfected with a GFP fusion to the indicated construct. The time is indicated in seconds. At time "0", addition of the Rho kinase inhibitor Y-27632 was added at 100 µM final concentration.

**Suppl. movie3.** TIRF analysis of VASP-mCherry expressing C2.7 cells co-transfected with a GFP fusion to the indicated construct. The time is indicated in seconds. At time "0", addition of the Rho kinase inhibitor Y-27632 was added at 100 µM final concentration.

**Suppl. Movie 4.** 1205Lu melanoma cells 1205Lu melanoma cells were transfected with the indicated constructs. Cells were perfused in a microfluidic chamber and allowed to adhere for 20 min prior to application of shear stress reaching 22.2 dynes.$cm^{-2}$. The elapsed time is indicated in seconds.
